# Linking Population Dynamics to Microbial Kinetics for Hybrid Modeling of Engineered Bioprocesses

**DOI:** 10.1101/2021.04.15.440059

**Authors:** Zhang Cheng, Shiyun Yao, Heyang Yuan

## Abstract

Mechanistic and data-driven models have been developed to provide predictive insights into the design and optimization of engineered bioprocesses. These two modeling strategies can be combined to form hybrid models to address the issues of parameter identifiability and prediction interpretability. Herein, we developed a novel and robust hybrid modeling strategy by incorporating microbial population dynamics into model construction. The hybrid model was constructed using bioelectrochemical systems (BES) as a platform system. We collected 77 samples from 13 publications, in which the BES were operated under diverse conditions, and performed holistic processing of the 16S rRNA amplicon sequencing data. Community analysis revealed core populations composed of putative electroactive taxa *Geobacter, Desulfovibrio, Pseudomonas*, and *Acinetobacter*. Primary Bayesian networks were trained with the core populations and environmental parameters, and directed Bayesian networks were trained by defining the operating parameters to improve the prediction interpretability. Both networks were validated with Bray-Curtis similarly, relative root-mean-square error (RMSE), and a null model. The hybrid model was developed by first building a three-population mechanistic component and subsequently feeding the estimated microbial kinetic parameters into network training. The hybrid model generated a simulated community that shared a Bray-Curtis similarity of 72% with the actual microbial community and an average relative RMSE of 7% for individual taxa. When examined with additional samples that were not included in network training, the hybrid model achieved accurate prediction of current production with a relative error-based RMSE of 0.8 and outperformed the data-driven models. The genomics-enabled hybrid modeling strategy represents a significant step toward robust simulation of a variety of engineered bioprocesses.

## 1. Introduction

Engineered bioprocesses are widely applied to treat waste streams and recover valuable resources (Rittmann and McCarty 2012). To facilitate the design and optimization of full-scale bioprocesses, a number of mechanistic and data-driven models have been developed over the past 60 years (Batstone et al. 2002, Bhat and McAvoy 1990, Henze et al. 2000).

Mechanistic models can provide predictive insights into the fundamental processes in biological systems (Jeppsson 1996). The model structure has been improved with a greater understanding of the microbiomes in biological systems (Donoso-Bravo et al. 2011, Henze et al. 2000, Ng and Kim 2007). For example, the Anaerobic Digestion Model No.1 composed of 19 bioconversion steps and 100 parameters is by far the most comprehensive mechanistic model formulated for engineered bioprocesses and can be readily modified to simulate specific applications (Liu et al. 2017, Rodríguez et al. 2008, Zhao et al. 2019). An inherent problem of this modeling strategy is that the model structure and parameters are largely unidentifiable (Donoso-Bravo et al. 2011, Jeppsson 1996). This stems from the simulation of microbial kinetics, in which the functional populations cannot be fully recapitulated by the Monod expressions, and the associated kinetic parameters cannot be directly measured (Donoso-Bravo et al. 2011, Ng and Kim 2007). Although kinetic parameters can be derived from biochemical measurements, they show considerable variations under different operating conditions (Bernard et al. 2006, Ng and Kim 2007). As a result, a mechanistic model developed for a specific bioprocess needs constant parameter calibration to cope with operational perturbations but still falls short when applied to other biological systems.

Data-driven models are not limited by identifiability issues and can yield more accurate predictions than mechanistic models do when a sufficiently large data pool is provided (Walpole et al. 2017). Artificial neural networks can be constructed with appropriate input variables and network architecture to predict the effluent quality (Mendes et al. 2015, Moral et al. 2008). Recent studies have demonstrated the applicability of several machine learning algorithms, including support vector machine, random forest, extreme gradient boosting, and k-nearest neighbors, to full-scale anaerobic digesters (De Clercq et al. 2019, Wang et al. 2020b). Despite the outstanding learning performance, most of the data-driven models are black boxes that are unable to generate interpretable predictions (Rudin 2019). This is particularly problematic for complex biological systems whose performance is largely determined by the microbial population and activity, and thus no mechanistic insights can be obtained from the simulation. Training data-driven models with microbial population dynamics presents a promising solution to tackle this issue. Previous studies have incorporated genomic data into machine learning models (neural networks and Bayesian networks) to reconstruct microbial communities in natural ecosystems (Kuang et al. 2016, Larsen et al. 2012). Using similar strategies, an array of data-driven models was constructed to simulate the performance and stability of engineered bioprocesses (Lesnik et al. 2020, Lesnik and Liu 2017, Yuan et al. 2017).

Hybrid models can potentially address the limitations of those two modeling strategies (Cote et al. 1995, Karama et al. 2001, Zhao et al. 1997). A common approach is to couple the mechanistic and statistical components in series. The error signals obtained from the mechanistic component are converted into those for the network component, which are subsequently used to update the network weights through back-propagation (Lee et al. 2002). Through such integration, hybrid models yield robust and semi-interpretable predictions for non-linear behaviors such as microbial kinetics (Zendehboudi et al. 2018). By far, all hybrid models are built with physical and biochemical parameters and thus unable to reveal the connections between microbial population and microbial kinetics. We propose to link them and improve the prediction robustness by incorporating genomic data into model construction.

The objectives of this study are to 1) comprehensively characterize the microbiome in a specific type of bioprocess and 2) formulate a novel hybrid model based on the population dynamics and microbial kinetics. To this end, we used bioelectrochemical systems (BES) as a platform system for model construction. As emerging biotechnology that simultaneously achieves water/wastewater treatment and energy/resource recovery (Logan et al. 2006, Wang and Ren 2013), BES are ideal for this task because they respond quickly to environmental perturbations (Yuan et al. 2016, Yuan et al. 2015), with current production acting as a sensitive indicator of the microbial population and functional dynamics. The microbial communities in BES are highly enriched with relatively low microbial diversity (Yates et al. 2012, Zhu et al. 2014), and hence represent a desirable level of complexity: diverse enough to be relevant to the microbiomes in other bioprocesses yet simple enough to be *in silico* reconstructed. To improve the compatibility of our models, we performed an extensive literature review and collected the 16S rRNA gene amplicon sequencing data from 77 samples in 13 publications, in which the BES were operated under a wide range of conditions. Core populations were selected at different taxonomic levels and used to train Bayesian networks, a machine learning model capable of characterizing the causal relationships among variables (Uusitalo 2007). Meanwhile, microbial kinetics were calculated using a three-population mechanistic component and fed into the training process to improve the prediction. This hybrid modeling strategy is expected to take advantage of the rapid growth of the genomic database (Kahn 2011), circumvent the time-consuming calibration of the kinetic parameters, and be broadly applicable to a variety of engineered bioprocesses.

## 2. Materials and Methods

### 2.1 Data collection and sequence processing

A comprehensive literature review was carried out, and 13 publications containing 77 samples were selected for downstream community analysis and model construction. The detailed information about the selected publications is listed in the Supporting Information (SI) Table S1. The selection criteria are: 1) 16s rRNA gene amplicon sequences were properly deposited in the National Center for Biotechnology Information (NCBI) or DNA Data Bank of Japan (DDBJ) databases for holistic sequence processing; 2) the results reported eight key parameters closely related to BES performance, including substrate composition and concentration, coulombic efficiency (CE), pH, current, anode area, external resistance, hydraulic retention time, and temperature; 3) the results included the variation of chemical oxygen demand (COD) over time or currents/voltage that can be used to calculate the time series of COD. The selected studies show a variety of reactor configurations, operation modes, substrates, and operating conditions (SI Table S1), which is expected to enhance the compatibility of the predictive models developed in the present study.

The sequence data from the selected publications contained both pair-end or single-end reads and were converted into a uniform format before further processing. Briefly, pair-end reads were merged using Vsearch (Rognes et al. 2016), and the chimeric and low-quality sequences were eliminated using the QIIME2 plug-in DADA2 (Callahan et al. 2016). The Greengenes database (gg_13_8 updated February 2011) was used to conduct sequence alignment and train the taxonomy classifier of the denoised sequences (DeSantis et al. 2006). In addition, due to the various primer-targeted regions (V1-V9) used to amplify 16s rRNA gene in those studies, the primer pair 8F/907R was set as the forward and reverse primers for the classifier to encompass all of the sequences from the samples.

### 2.2 Community analysis

Alpha (Shannon and Simpson indices) diversity analysis, principal coordinate analysis (PCoA), and redundancy analysis (RDA) were performed using R to unravel intra-sample diversity and inter-sample distance. Core populations were selected at the genus, order, and phylum levels with the following criteria (Yuan et al. 2019): 1) at least one occurrence in the 13 studies with the relative abundance ≥0.05% and 2) average relative abundance ≥2% across all 77 samples. The criteria allow us to retain the major functional populations in the microbial community and compress the genomic data for downstream model construction. For this reason, core population was not selected at the operational taxonomic unit (OTU) level as the 100%-similarity clustering strategy generated over 9,000 OTUs, and the majority of the samples have little overlap on the community composition. To build a phylogenetic tree for the core genera, the sequence of the most abundant OTU within a core genus was selected as a representative. The phylogenetic tree was built using ARB (Ludwig et al. 2004), and the *Silva* database (LTPs132_SSU.arb for 16s rDNA updated June 2018) was used for sequence alignment (Quast et al. 2013).

### 2.3 Bayesian network analysis

To prepare for network construction, the abundance of the core taxa and the values of the environmental parameters were scaled to 0 – 1 (Bishop 2013):

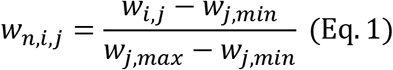

where *w*_*i,j*_ is the relative abundance of population *j* in sample *i*, *w*_*j,max*_ is the maximum relative abundance of population *j*, *w*_*j,min*_ is the minimum relative abundance of population *j*, and *w*_*n,i,j*_ is the normalized abundance of population *j* in sample *i*. Structure learning and parameter learning were performed using the hill-climbing algorithm and maximum likelihood estimation, respectively (Scutari 2010).

Primary networks (SI Figures S4A, S5A, and S6A) were trained without considering the *a priori* knowledge about the operating conditions, and the node directions were solely inferred by the network algorithm. Directed networks (SI Figures S4B, S5B, and S6B) were trained by defining temperature, anode electrode area, external resistance, and hydraulic retention time as the parent nodes. This was because those parameters remained unchanged throughout the operation and thus were not affected by other parameters. A blacklist function was applied to define the unidirectional relationships between those operating parameters and other variables. The same training procedures were conducted at the genus (38 core taxa), order (32 core taxa), and phylum (13 core taxa) levels. Considering the sample quantity and computational cost, a leave-one-out cross-validation strategy was selected for three validation methods (Bro et al. 2008): Bray-Curtis similarity between the predicted and observed microbial community, relative root-mean-square error (RMSE, Eq. 2), and null model analysis.

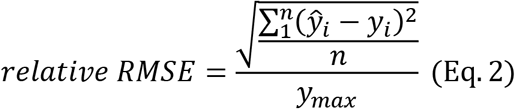

where *ŷ_i_* is the predicted value; *y*_*i*_ is the observed value; *y*_*max*_ is the maximum observed value; and *n* is the number of samples. Null models were constructed by setting the abundance of all core taxa to the average abundance across all samples (Gotelli 2002).

### >2.4 Hybrid model construction

A hybrid model was constructed at the genus level to bridge population dynamics and microbial kinetics following the procedures shown in SI Figure S1. Briefly, maximum substrate utilization rates, maximum growth rates, and mediator yield were calculated using the mechanistic component described below and included as the nodes for network training. Although these kinetic parameters are unmeasurable, the normalization step (Eq. 1) converts exact values into a general tendency of microbial activity that can be statistically connected to the actual relative abundance of the core population (Weissman et al. 2021), and thus the estimated kinetic parameters do not need to be validated.

The mechanistic component assumes two-step degradation (SI Figure S2) of the substrates by three populations (Pinto et al. 2011). At the first stage, complex organic matters such as polysaccharides, lipids, and proteins are decomposed by primary degraders to low-molecular intermediate products including acetic acid, propionic acid, ethanol, etc. At the second stage, electroactive and non-electroactive microbes convert the intermediates into electrical energy and methane, respectively. Take electroactive microbes as an example, the mass balance for the growth and activity are described using Eq. 3 and Eq. 4, respective:

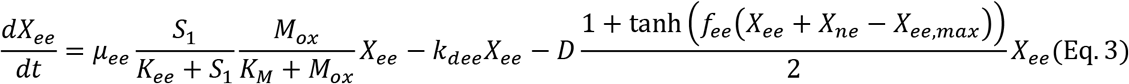

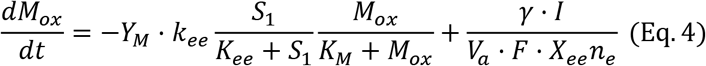

where *X*_*ee*_ and *M*_*ox*_ are the concentrations of electroactive microbes and mediator, respectively; *K*_*ee*_ and *K*_*M*_ are the substrate half-saturation constant and mediator half-saturation constant, respectively; *μ*_*ee*_ and *k*_*d,ee*_ are the growth and decay rates, *X*_*ee,max*_ is the maximum capacity of electroactive microbes in the anode; *Y*_*M*_ is the mediator yield; *γ* is the molecular mass of mediator; *I* is the current production; *V*_*a*_ is the anode volume; *F* is the Faraday constant (A/d·mol); and *n*_*e*_ is the number of electrons transfer. The mass balance was modified based on the multiplicative Monod expressions from previous studies (Ping et al. 2014, Pinto et al. 2010). The detailed formulation of the mechanistic component and the parameters can be found in the SI Method and Table S2. Because the selected publications provided multiple types of data and operated the reactors under distinct conditions, the mechanistic component was slightly modified for individual reactors (SI Table S3), and Literature #27 (S27) was not considered for mechanistic modeling because a time series of COD was not available.

The maximum substrate utilization rate and maximum growth rate are considered to be the most critical values for simulation of engineered bioprocesses (Rittmann and McCarty 2012), and mediator yield is a unique parameter for BES. Those kinetic parameters were estimated with specific limits according to previous studies while others parameters were retrieved from literature (Kato Marcus et al. 2007, Wilson and Kim 2016). To estimate the parameters, the total substrate concentration over time and the initial values of the kinetic parameters were collected, calculated, or estimated from the selected papers. Because biomass and mediator concentrations were not measurable and unavailable in the literature, some assumptions are made: 1) primary degraders have an initial concentration of 100 mg/L in all reactors, 2) electroactive and non-electroactive microbe grow evenly on the anode surface at an average thickness of 60 μm (Lee et al. 2009, Torres et al. 2008), 3) the maximum biofilm capacity in BES was assumed to be 600 mg/L (Ping et al. 2014), and 4) the electroactive microbe fraction is proportional to the CE. For microbial electrolysis cells whose CE was higher than 100% because of the applied voltage, the concentration of electroactive microbes was assumed to be 500 mg/L. Because the dimension of the anode electrode was not directly provided in some of the studies, the area was estimated based on the specific area and size of the electrode (Logan et al. 2007).

### 2.5 Model prediction

Six additional publications were collected to further demonstrate the robustness of the hybrid model (SI Table S7). In the first step, the operating parameters (i.e., temperature, anode electrode area, external resistance, and hydraulic retention time) and pH from those studies were input into the Bayesian networks to predict current and CE directly. In the second step, the kinetic parameters obtained from the Bayesian network were input into the mechanistic component to calculate the steady-state COD, which was then converted into current production based on the expression of CE (Logan et al. 2006). The predicted and observed values were compared using an RMSE based on relative errors (Walpole et al. 2017):

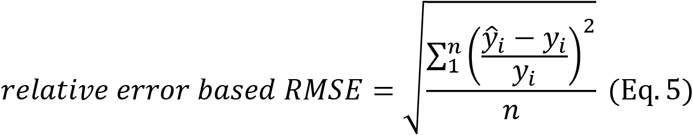

## 3. Results and Discussion

### 3.1 BES operated under a variety of conditions

The 13 selected studies contained 39 samples from microbial fuel cells, 31 samples from microbial electrolysis cells with external voltage input, and 7 samples from microbial desalination cells with saline environments (SI Table S1). In addition to different reactor configurations, the bioreactors were fed with a variety of substrates whose main organic matter could be categorized as non-fermentable (acetate and methanol), fermentable (glucose, ethanol, and propyl alcohol), or complex (brewery wastewater, food waste, and pig slurry). The selected studies also presented multiple operation modes including batch, continuous, and continuous with pulse substrate loading. In terms of the performance, the 77 samples showed significant difference (SI Table S4) in CE which ranged from 0.02% (due to high external resistance, e.g., S22) to 140% (due to applied voltage, e.g., S37). Overall, the selected samples have included the conditions commonly found in BES studies and are expected to yield representative core populations and models.

### 3.2 Core population in BES

The 2.6 million sequence reads from the 77 samples resulted in approximately 9600 OTUs, based on which the alpha diversity was analyzed. Although the diversity indices (Shannon, Simpson, and Chao1, SI Table S5) did not follow a normal distribution as revealed by the Shapiro-Wilk analysis results (p < 0.05), it could well reflect the variation in operating conditions. For example, the Shannon and Simpson indices for the samples fed with complex substrates (i.e., S15, S17, S20, S27, and S42) were 4.25 and 0.96, respectively, significantly higher than the 2.24 and 0.73 of the samples fed with non-fermentable substrates. Similar results were reported in previous studies (Wang et al. 2020a), and a highly diverse microbial community was expected to enhance system stability (Girvan et al. 2005). In addition, the majority of the samples fed with non-fermentable and fermentable substrates showed a Chao1 index of approximately 100, indicating that those BESs were sufficiently sampled, and the key microbes could be captured when selecting the core population. This is confirmed by the rarefaction curves presented in some of the selected publications.

PCoA based on Bray-Curtis distance showed a critical role of substrate composition in microbial community assembly (Figure 1). Specifically, samples cultivated with starch- and yeast extraction-based synthetic wastewater (S20 and S22) were found in the top right corner of the PCoA graph, while those with complex food waste, brewery wastewater, and pig slurry (S15, S17, S42, and S51) were clustered in the center. S48 used ethanol as the sole carbon source and was isolated from other samples. Additionally, the anode area appeared to be an important factor that drove the microbial community assembly in S48 (SI Figure S3), which was amended with granular activated carbon in the anode. Another key deterministic factor of microbial community assembly is the seed source. Unlike other studies, S49 was inoculated with activated sludge and formed a distinct community structure. Similarly, activated sludge was the seed of S35 and together with temperature (SI Figure S3) led to communities significantly different from other studies. In summary, varied substrate composition, reactor configuration, and operation mode provided a comprehensive pool of microbial communities for model construction.

**Figure 1.**
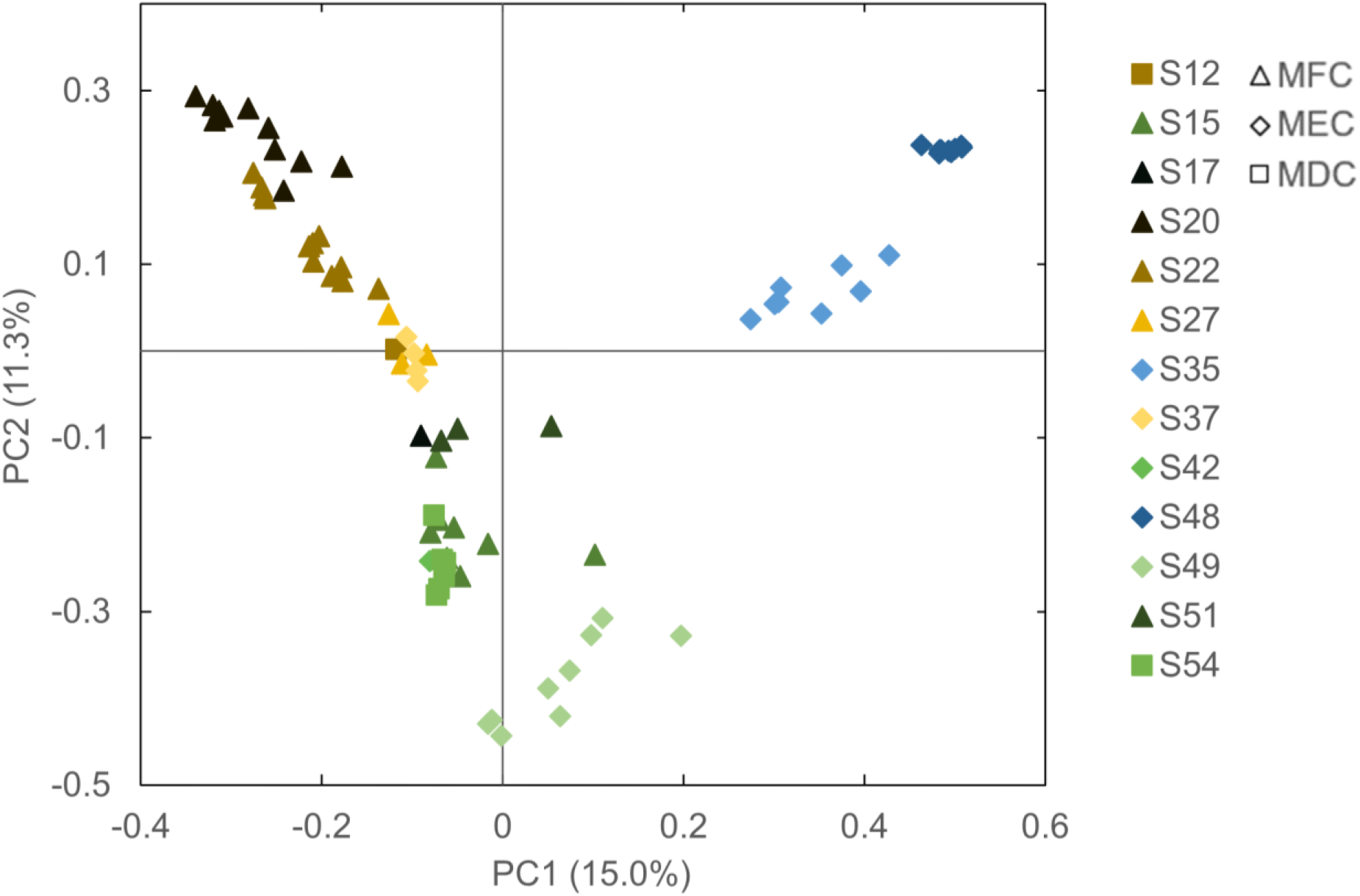
Bray-Curtis distance-based PCoA of the 13 selected studies. MFC: microbial fuel cells, MEC: microbial electrolysis cells, MDC: microbial desalination cells.

Core populations were selected at different taxonomic levels based on the occurrence (at least one occurrence in the 13 studies with the abundance ≥0.05%) and abundance (≥2% across all 77 samples) (Ling et al. 2016, Saunders et al. 2016). At the genus level, 38 core taxa were identified, accounting for 55% of the abundance on average. The selection criteria were considered stringent given that the bioreactors were operated under distinct conditions, and the microbial communities were highly diverse. This was reflected by the loss of several abundant taxa in specific BES. For instance, the core genera made up of less than 20% of the abundance in some of the samples due to the unique flow pattern (plug-flow, S22), substrate (propyl alcohol, S27), and reactor configuration (applied voltage, S37). Nonetheless, the core population included some well-characterized genera such as *Geobacter*, *Desulfovibrio*, *Pseudomonas*, and *Acinetobacter*, which were frequently found abundant in BES and potentially involved in current production.

The presence of *Geobacter* often serves as the indicator to explain the BES performance, in particular current production, because a few members of this genus are highly efficient in extracellular electron transfer (EET) (Logan et al. 2019, Lovley et al. 2011). Indeed, *Geobacter* was identified to be a core taxon (G29, Figure 2) with an average abundance of 15% across all samples and an individual abundance higher than 2% in 39 samples. This genus was dominant in S20 (17%), S35 (71%), S48 (27%), and some of the samples in S15 (9%) and S49 (8%), which all showed a CE over 15%. However, *Geobacter* was not present in other high-CE samples likely because the operating conditions (e.g., high salinity in S12 and S54) did not favor its growth (Miyahara et al. 2015)*. Desulfovibrio* spp. from the same class of Deltaproteobacteria are common sulfate-reducing bacteria whose EET ability has also been reported (Aulenta et al. 2012, Gacitúa et al. 2014, Yu et al. 2011). This genus (G28, Figure 2) was abundant in 13 samples (>2%) and dominant in ethanol-fed S48 (27%). *Desulfovibrio* is known to oxidize ethanol with sulfate as the electron acceptor. In the absence of sulfate, *Desulfovibrio* can still grow syntrophically with methanogens and oxidize the ethanol to acetate through interspecies hydrogen transfer (Hensgens et al. 1993, Kremer et al. 1988).

**Figure 2.**
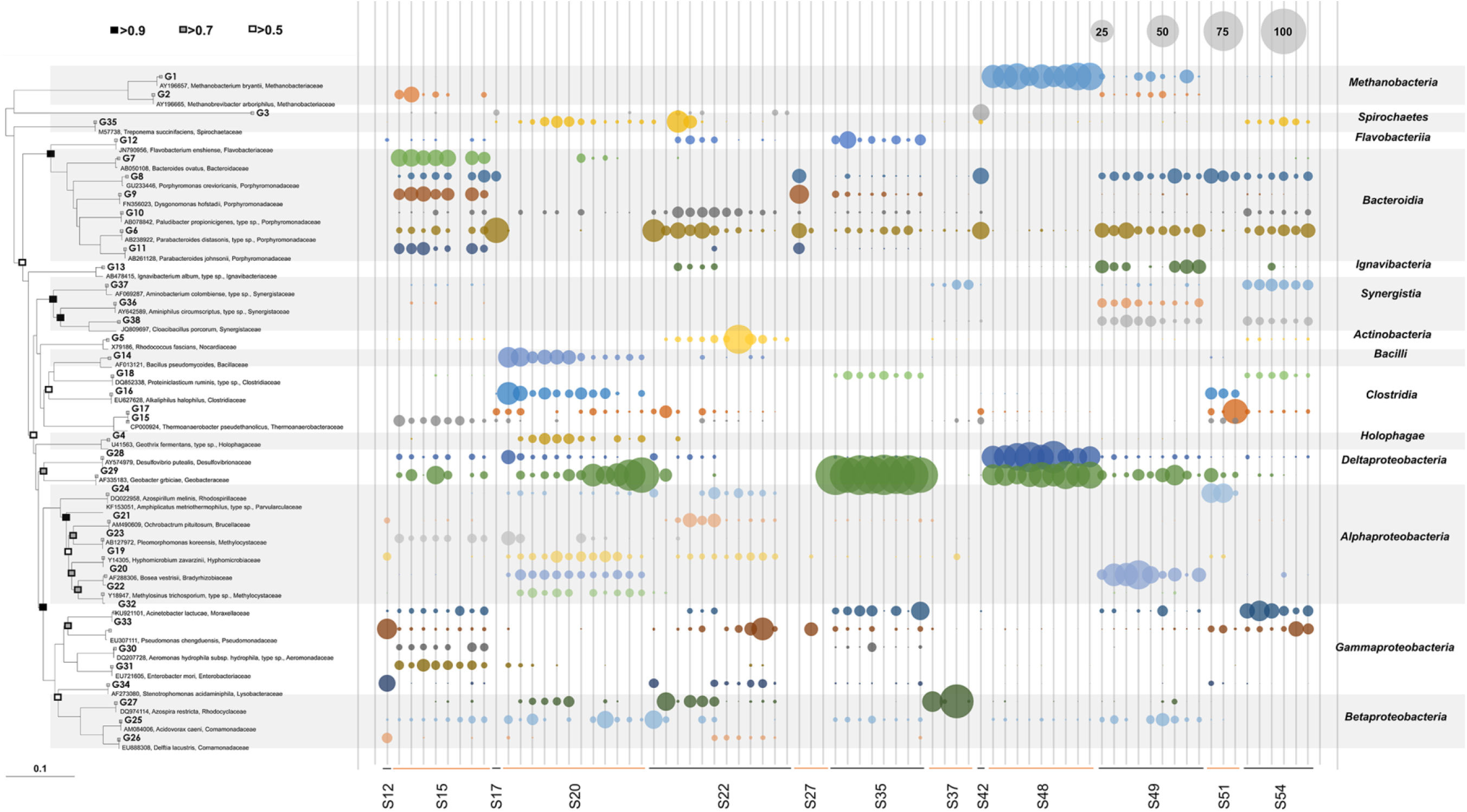
Phylogenetic tree and relative abundance of 38 core genera selected from the 77 samples in 13 publications.

*Pseudomonas*, a well-studied genus forming biofilm in many anaerobic environments, is another core taxon that has been reported to carry out EET by using phenazines as an electron shuttle (Rabaey et al. 2004). *Pseudomonas* (G33, Figure 2) was widely present in S12, S15, S22 S35, S49, and S54. This genus potentially plays a critical role in shaping the microbial community structure as the phenazines actively produced for quorum sensing can be scavenged for electron shuttling by other species such as *Acinetobacter*. As shown in Figure 2, *Acinetobacter* (G32) was found in samples where *Pseudomonas* was abundant and was previously speculated to utilize phenazines as electron shuttles for EET (Liu et al. 2013, Yuan et al. 2017). Because EET via electron shuttles is limited by diffusion and the conductivity of the anolyte (Torres et al. 2010), *Pseudomonas* and its metabolic partners are less ubiquitous than *Geobacter* in BES, and *Pseudomonas*-dominated communities are more likely to be found in specific environments such as microbial desalination cells with a high ion concentration (Yuan et al. 2017).

Several genera from the class Bacteroidia were found to be abundant (Figure 2). Previous studies suggested that this group of bacteria could degrade complex organic compounds including proteins, polysaccharides, and pectins (Dongowski et al. 2000, Grenier et al. 1989). For instance, *Parabacteroides* (G6 & G11, Figure 2) show an abundance higher than 5% in most of the samples fed with complex organic. It has also been reported that some members in the class Bacteroidia degrade biomass and can serve as scavengers of dead cells (Madigan 2014, Reichenbach 1992). Those taxa might act as degraders of soluble organic matter and provide electroactive microbes simple substrates (Tan et al. 2012, Zeppilli et al. 2020). In addition to fermentative bacteria, methanogens were observed with considerable abundance in S48 (Figure 2) and possibly carried out syntrophic electron transfer with ethanol-consuming in the presence of conductive granular activated carbon (Yuan et al. 2018).

Overall, microbial community analysis demonstrated a core BES population composed of primary fermentative bacteria that convert complex organic matter to simple electron donors, and electroactive microbes and their competitors (e.g., methanogens) growing on the fermentative products. The results thus justify the modeling of BES based on those three guilds (Pinto et al. 2011). However, such a model structure is incapable of differentiating the contribution of different types of electroactive microbes and their EET mechanisms (i.e., direct contact vs. electron shuttling) due to the experimental challenge to measure the associated biochemical parameters (e.g., the concentration of phenazines and other electron shuttles). The same pitfall is also found in mechanistic modeling of activated sludge and anaerobic digestion, in which the core populations consist of functionally redundant taxa occupying the same ecological niches (Ju and Zhang 2015). This explains the constant parameter calibration required by mechanistic models. To address the issue and improve the prediction robustness, a new modeling approach is imperative.

### 3.3 Reconstruction of microbial community

Two types of Bayesian networks were trained with the same dataset containing environmental parameters and relative abundance of the core populations (SI Figure S4-S6): primary networks whose node directions were not restricted by *a priori* knowledge and directed networks in which the operating parameters were set as the parent nodes. The latter were constructed based on the fact that operating parameters such as external resistance and hydraulic retention time remained unchanged throughout the operation and hence should be not affected by microbial community dynamics and system performance.

To validate the modeling approach and evaluate the prediction of the community structure, Bray-Curtis similarity between the predicted and observed communities was calculated. At the genus level, the directed network achieved the most accurate prediction, followed by the primary network and a null model (Bray-Curtis similarity: 0.72 > 0.64 > 0.52, p < 0.05, SI Figure S7A). Similar trends were observed at the order and phylum levels, but the prediction accuracy did not show consistent improvement as the taxonomic level increased. The Bray-Curtis similarity from the directed networks dropped slightly to 0.61 at the order level and raised back to 0.79 at the phylum level. The results were not in agreement with the previous findings that prediction accuracy could be continuously improved by training data-driven models at higher taxonomic levels (Kuang et al. 2016, Yuan et al. 2017). It should be noted that those models were built based on highly specific environments and communities (e.g., acid mine drainage and microbial desalination cells), whereas the models in the present study considered a variety of environmental conditions, which might statistically compromise the model robustness (Walpole et al. 2017).

To further validate the modeling approach, relative RMSE was calculated for the 6 environmental parameters and 38 core genera (SI Figure S8-S10). At the genus level, the RMSE values from the directed and primary network were 2% - 17% and 3% - 24%, respectively. The abundances of putative electroactive taxa *Desulfovibrio*, *Pseudomonas*, and *Acinetobacter* were well estimated by both networks with the RMSE ranging from 4% to 10%. On the other hand, the RMSE for *Geobacter* was improved from 24% with the primary network to 16% with the directed network. The poor prediction of *Geobacter* is likely because some members from this genus, despite their dominance in many anaerobic environments (Lee et al. 2016, Lin et al. 2017), are inefficient in or incapable of EET (Lovley et al. 2011, Rotaru et al. 2015). Similar to Bray-Curtis similarity, RMSE was improved at the phylum but not at the order level (SI Figure S9 and S10). The phylum Proteobacteria, which includes the putative electroactive taxa discussed above, is estimated with the highest accuracy (relative RMSE <1%). Overall, the more accurate prediction from the directed networks at all three taxonomic levels suggests that the modeling approach can be enhanced by introducing reasonable structure control.

After the modeling approach was validated with Bray-Curtis similarity and RMSE, final networks were constructed from the whole dataset to infer microbial interactions (SI Figure S4-S6). In the genus-level networks, putative electroactive taxa *Geobacter* (G29), *Desulfovibrio* (G28), and *Pseudomonas* (G33) did not show any association with CE and current, whilst *Acinetobacter* (G32) was not correlated with the system output. The networks at higher taxonomic levels yielded even less interpretable inference regarding the potential functions of the core taxa. For example, methanogens were predicted to be more related to current production than Proteobacteria (SI Figure S6A). The results collectively indicate that more *a priori* knowledge needs to be included in model training to improve the robustness and interpretability of the inference.

### 3.4 Hybrid modeling of BES performance

To build a hybrid model, the rates for substrate utilization and microbial growth and mediator yield were first estimated using the three-population mechanistic component (SI Table S4). Some of the estimates were constant during calibration (e.g., 15 /d for substrate utilization rate) because they were the boundary values determined based on previous studies (Bruce and Perry 2001, Kato Marcus et al. 2007, Wilson and Kim 2016). The estimated microbial kinetic parameters were subsequently included in the training dataset to construct a hybrid network at the genus level (Figure 3). It should be noted that the scaled kinetic parameters represent the trend of microbial activity and thus do not require accurate estimation or validation. A whitelist function was further applied to force putative electroactive genera *Geobacter, Desulfovibrio, Pseudomonas,* and *Acinetobacter*to to directly affect current generation and improve the prediction interpretability. The hybrid network yielded a simulated community that shared a Bray-Curtis similarity of 0.72 with the actual genera-level core population, which was comparable to the result from the directed network and significantly better than that of the null model (t-test, p < 0.05). The relative RMSE of the hybrid model ranging from 3% to 18% was also similar to those from the directed network.

**Figure 3.**
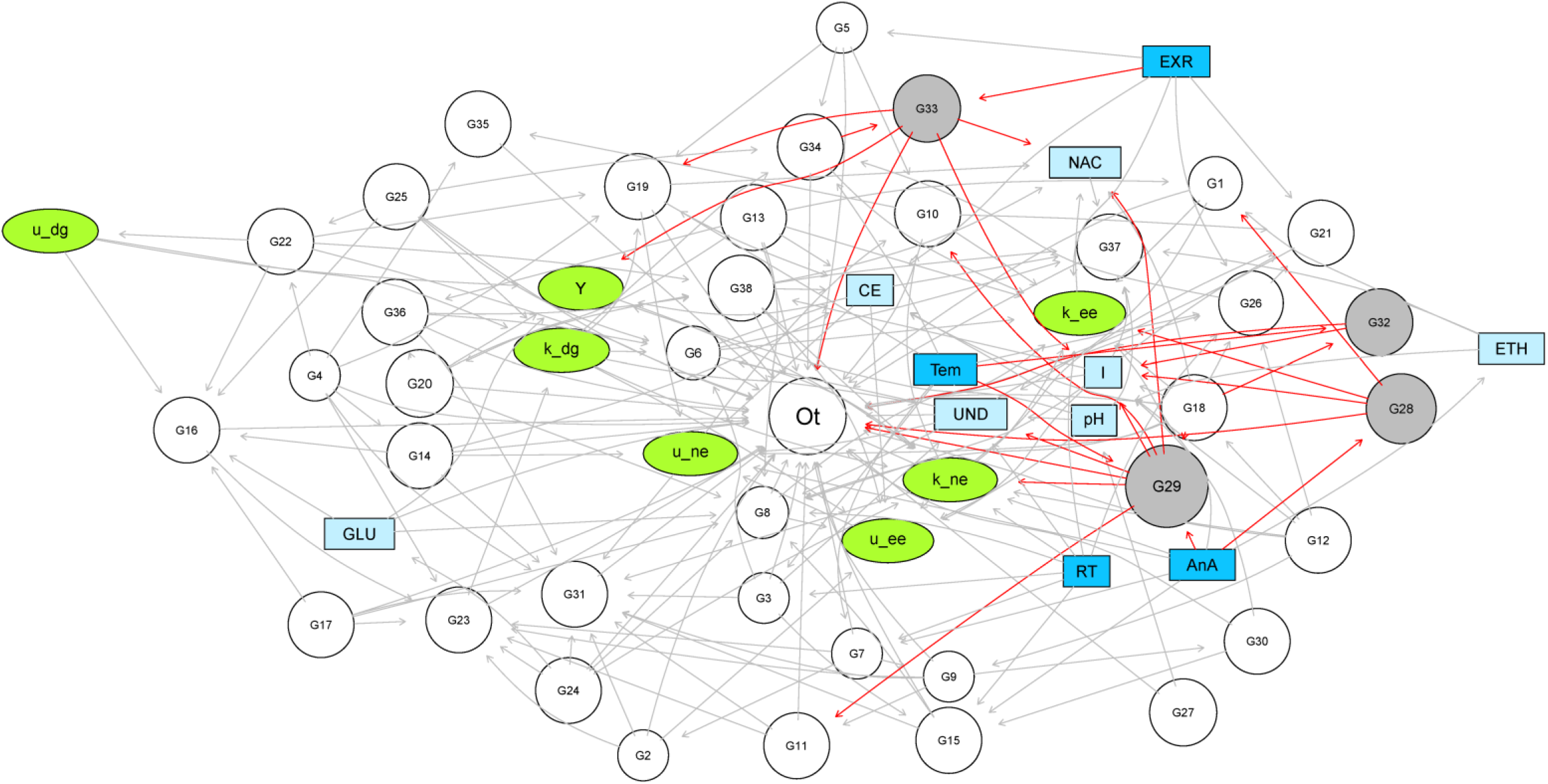
The hybrid Bayesian network at the genus level. *Geobacter* (G29)*, Desulfovibrio* (G28)*, Pseudomonas* (G33), and *Acinetobacter* (G32) are highlighted in grey and forced to directly affect current. The symbols for the core genera can be found in SI Table S6. Light blue nodes are biochemical parameters. Dark blue nodes are operating parameters unaffected by other nodes and serve only as the parent nodes. Green nodes are kinetic parameters estimated using the mechanistic component, in which u_dg, u_ee, and u_ne are the maximum growth rates for primary degraders, electroactive microbes, and non-electroactive microbes, respectively. k is the maximum substrate utilization rate and Y is the mediator yield. EXR: external resistance, NAC: acetate, CE: coulombic efficiency, I: current, Tem: temperature, ETH: ethanol, GLU: glucose, RT: hydraulic retention time, AnA: anode area, UND: undefined substrate.

The hybrid network generated reasonable inference of the relationships between microbial population and kinetics (Figure 3), as evidenced by the strong positive correlation of the EET-related substrate utilization rate with *Desulfovibrio* (coefficient = 0.86), as well as the positive correlation of mediator yield with *Pseudomonas*. Meanwhile, mediator yield was negatively related to glucose, likely because fermentable substrates could lead to significant electron loss (Parameswaran et al. 2010). The EET-related substrate utilization and growth rates were both associated with the genera (G6 & G8) from the class Bacteroidia. As discussed above, those taxa can degrade dead cells and soluble microbial products, thereby creating a favorable environment for electroactive microbes (Ni et al. 2011, Ni et al. 2010). Despite those biologically sound inferences, the hybrid model still contained unexplainable interactions such as the negative association between the EET-related substrate utilization rate and anode area, underpinning the elimination of data-driven models in prediction interpretability.

The developed models were examined with six new samples that were not included in network training, and the hybrid model (hybrid network + mechanistic component) achieved the most accurate prediction of current production compared with the data-driven models. It can be seen from Figure 4 that the predicted results from the hybrid model agree well with the experimental values with slight deviation at the high current range. The low relative error-based RMSE of 0.8 further indicates outstanding prediction accuracy throughout the examined current range. The hybrid network alone loses the prediction power at high current, resulting in a higher relative error-based RMSE of 6.7, whereas the directed network is incapable of generating satisfactory prediction and shows the highest relative error-based RMSE of 16.3. The significantly improved prediction performance of the hybrid model likely stems from the close connection between population dynamics and microbial kinetics. Under a given condition, each population (either a single species or a functional guild) has specific maximum substrate utilization and growth rates that are largely determined by its unique lifestyle and ecophysiology (Rittmann and McCarty 2012), which can thus be statistically inferred from the genomic data (Weissman et al. 2021). On the other hand, system performance such as current production is affected by not only microbial population and activity, but also many other operating parameters including electrolyte conductivity and external resistance. Data-driven models that infer system performance directly from the microbial population do not consider the contribution of those operating parameters and hence cannot consistently yield accurate predictions.

**Figure 4.**
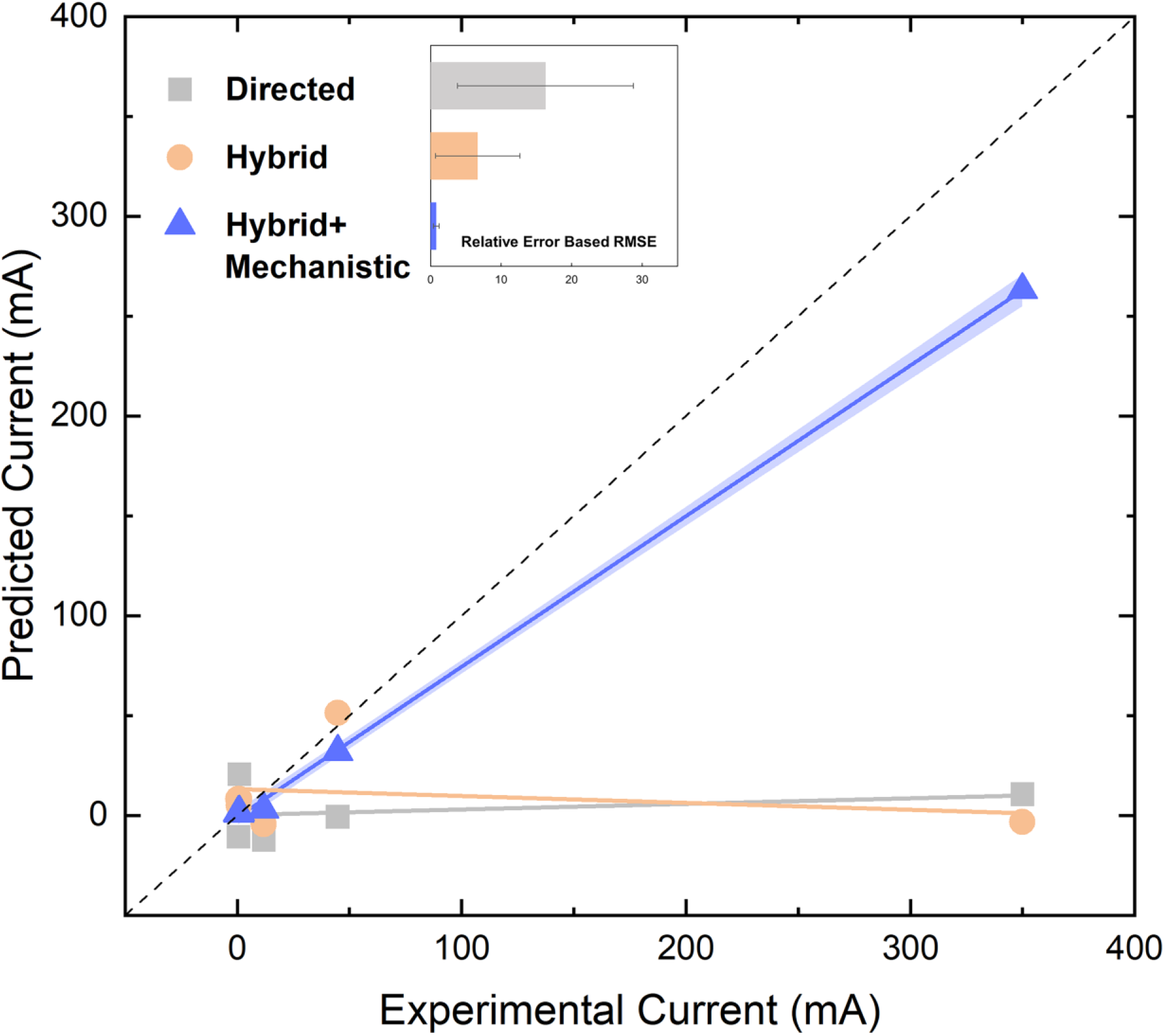
Comparison of experimental and predicted current from the directed Bayesian network, hybrid Bayesian network, and hybrid model (hybrid network + mechanistic component). Inset in the relative error-based RMSE.

Despite the robust performance, the hybrid model is not ready for practical implementation as accurate prediction can only be obtained with microbial community information as the input. When the data-driven component is fed solely with operating parameters, and the simulated microbial community serves as the intermediate to estimate the kinetic parameters, the prediction error quickly builds up along the inference, causing considerable uncertainty to the final prediction. Another challenge is that inadequate biochemical and sequencing data from the selected publications compromise the compatibility of both the data-driven and mechanistic components. These issues will be addressed in future studies with proper experimental design and alternative machine learning algorithms such as neural networks and random forest. Ultimately, the hybrid modeling approach is expected to be broadly applicable to various engineered bioprocesses including anaerobic digesters, activated sludge processes, anaerobic ammonium oxidation, etc.

## 4. Conclusion

We collected 77 samples from 13 studies in which the BES were operated under diverse conditions. Community analysis revealed a core population composed of primary fermentative bacteria, putative electroactive taxa *Geobacter, Desulfovibrio, Pseudomonas,* and *Acinetobacter*, as well as non-electroactive microbes such as methanogens. Bayesian networks were trained with the core populations and validated with Bray-Curtis similarity, relative RMSE, and a null model, all based on a leave-one-out cross-validation strategy. A hybrid model was built by combining mechanistic modeling and network training and achieved more accurate prediction of current production than data-driven models. This study provides insights into incorporating genomic data into hybrid modeling for robust and interpretable prediction.

## Acknowledgement

This work was supported by the U.S. Department of Agriculture [Award No. 2020-67019-31027].

