## Supporting information for "Linking Population Dynamics to Microbial Kinetics for Hybrid Modeling of Engineered Bioprocesses"

Intended for ***Water Research***

Type of contribution***: Research Article***

* **Corresponding author**

;

**Materials and Methods**

**Mass balance for primary degraders**

$$\frac{{dS}_{0}}{dt}=D\left( S_{0,in}-S_{0} \right)-k_{p}\frac{S_{0}}{K_{p}+S_{0}}X_{p} (Eq. 1)$$

$$\frac{{dX}_{p}}{dt}=\mu_{p}\frac{S_{0}}{K_{p}+S_{0}}X_{p}-k_{dp}X_{p}-D{\cdot X}_{p} (Eq. 2)$$

where $S_{0}$ is the concentration of complex substrates (mg/L); $S_{0,in}$ is the concentration of complex substrate in the influent (mg/L); $X_{p}$ is the concentration of the primary degraders (mg/L); $k_{p}$ is the maximum complex substrate utilization rate of the primary degraders (/d);$\mu_{p}$ is the growth rate of the primary degraders (/d); $K_{p}$ is the complex substrate half-saturation constant for the primary degraders (mg/L); $k_{dp}$ is the decay rate of the primary degraders (/d); and $D$ is the dilution rate (/d).

**Mass balance for electroactive and non-electroactive microbes**

$$\frac{{dS}_{1}}{dt}=D\left( S_{1,in}-S_{1} \right)+f_{e}\cdot k_{p}\frac{S_{0}}{K_{p}+S_{0}}X_{p}-k_{ee}\frac{S_{1}}{K_{ee}+S_{1}}\frac{M_{ox}}{K_{M}+M_{ox}}X_{ee}-k_{ne}\frac{S_{1}}{K_{ne}+S_{1}}X_{ne} (Eq. 3)$$

$$\frac{{dX}_{ee}}{dt}=\mu_{ee}\frac{S_{1}}{K_{ee}+S_{1}}\frac{M_{ox}}{K_{M}+M_{ox}}X_{ee}-k_{dee}X_{ee}-D\frac{1+\tanh\left( f_{ee}\left( X_{ee}+X_{ne}-X_{ee,max} \right) \right)}{2}X_{ee}(Eq. 4)$$

$$\frac{{dX}_{ne}}{dt}=\mu_{ne} \frac{S_{1}}{K_{ne}+S_{1}}X_{ne}-k_{dne}X_{ne}-D\frac{1+\tanh\left( f_{ne}\left( X_{ee}+X_{ne}-X_{ne,max} \right) \right)}{2}X_{ne} (Eq. 5)$$

$$\frac{{dM}_{ox}}{dt}={-Y}_{M}\cdot k_{ee}\frac{S_{1}}{K_{ee}+S_{1}}\frac{M_{ox}}{K_{M}+M_{ox}}+\frac{\gamma\cdot I}{V_{a}\cdot F\cdot X_{ee}n_{e}} (Eq. 6)$$

where $S_{1}$ is the concentration of simple substrates (mg/L); $S_{1,in}$ is the concentration of simple substrate in the influent (mg/L); $X_{ee}$, $X_{ne}$, and $M_{ox}$ are the concentrations of the electroactive microbes and non-electroactive microbes (mg/L), and mediator (dimensionless), respectively; $k_{ee}$ and $k_{ne}$ are the maximum substrate utilization rate of the electroactive microbes and non-electroactive microbes (/d), respectively;$\mu_{ee}$ and $\mu_{ne}$ are the growth rate of the electroactive microbes and non-electroactive microbes (/d), respectively; $K_{ee}$, $K_{ne}$, and $K_{M}$ are simple substrate half-saturation constants for the electroactive microbes, non-electroactive microbes (mg/L), and mediator half-saturation constant for the electroactive microbes (dimensionless), respectively; $k_{d,ee}$ and $k_{d,ne}$ are the decay rate of the electroactive microbes and non-electroactive microbes (/d), $X_{ee,max}$ and $X_{ne,max}$ are the maximum capacity of electroactive microbes and non-electroactive microbes in the anode; $f_{e}$ is the fraction of complex degradation for energy, 0.78 (Bruce and Perry 2001); $Y_{M}$ is the mediator yield for electroactive microbes; $\gamma$ is the molecular mass of mediator (mg/mol); $I$ is the current trough the circuit of BES (A); $V_{a}$ is the anode volume (L); $F$ stands for Faraday constant (A/d·mol); and $n_{e}$ represent the number of electrons transfer when a mediator is used by electroactive microbes.

**Equations trained by hybrid model for BES performance and microbial kinetic parameter prediction**

CE=0.173391763-0.559777461*unknown Bacteroidales+0.316382607*unknown Ignavibacteriaceae-0.206926425*Bacillus+0.392747536*Proteiniclasticum-0.257766*unknown Rhizobiales+0.261935899*vadinCA02+0.813563239*pH-0.252401161*EXR

I=0.077072495-0.417036918*Flavobacterium+0.33610861*Proteiniclasticum-0.005635669*Desulfovibrio-0.200705351*Geobacter-0.147272441*unknown Aeromonadaceae+0.670252023*Acinetobacter-0.033374272*Pseudomonas

u_dg=0.019635288+0.398282116*Methylosinus

k_dg=-0.008522951+0.751327026*Pleomorphomonas+0.3140015*unknown Comamonadaceae+0.847411887*u_dg

Y=0.165506276+0.558821608*Bosea+0.982721599*Pseudomonas+1.244204366*CE-0.608173418*GLU-0.636717071*pH+0.847542855*u_dg

k_ne=0.543862406+0.748636753*unknown Bacteroidales+0.517999463*Flavobacterium-0.324123009*Azospirillum+0.386043978*Geobacter-0.53628668*vadinCA02-0.516108345*AnA-0.424076894*RT-0.524581387*k_dg-0.192222373*Y

u_ee=0.460204837+0.470018028*Methanobrevibacter+0.367325204*unknown Porphyromonadaceae+0.367949836*vadinCA02+0.882608846*CE-0.448077724*pH-1.096109854*I-0.259567559*k_dg-0.277747232*k_ne

u_ne=0.481391764-0.290031466*Bacillus-0.163367299*Proteiniclasticum-0.307298133*unknown Rhizobiales-0.381517107*CE-0.402115684*UD-0.229567444*AnA-0.255466235*u_ee

k_ee=-0.121644971+2.145202816*Methanobacterium+0.563831876*unknown Bacteroidales-0.344508764*unknown Porphyromonadaceae+0.553617992*Paludibacter-0.250685831*unknown Ignavibacteriaceae+0.588642112*Proteiniclasticum+0.857639383*Desulfovibrio+0.85720023*Stenotrophomonas-1.861615332*AnA+0.674823585*EXR+0.949005788*RT+0.372685518*k_ne

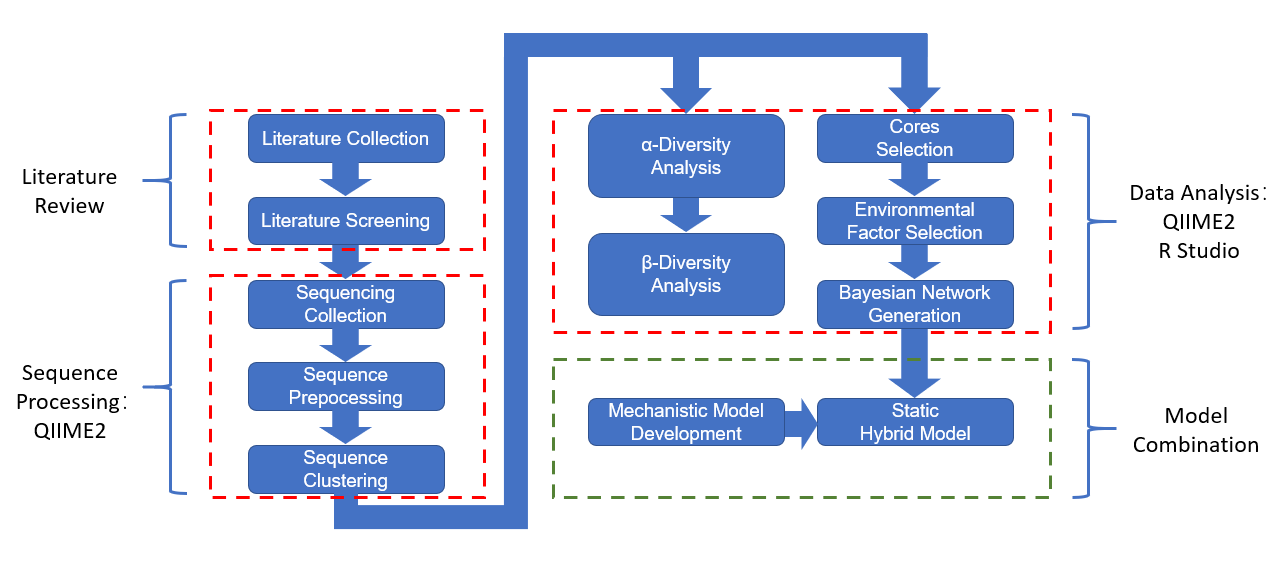

Figure S1. Overview of the hybrid modeling strategy.

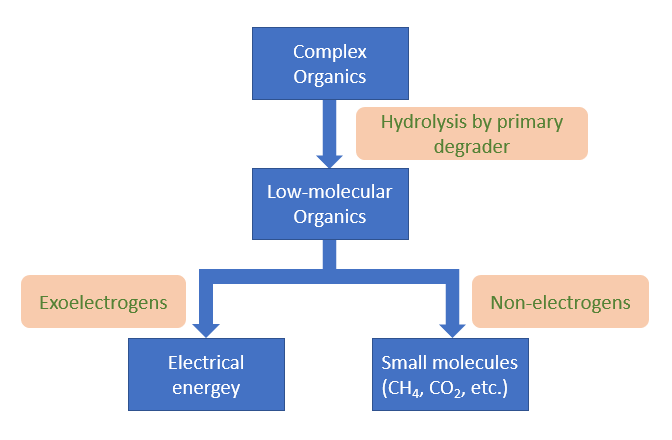

Figure S2. The presumed two-step degradation of complex organics by primary degraders, electroactive microbes, and non-electroactive microbes.

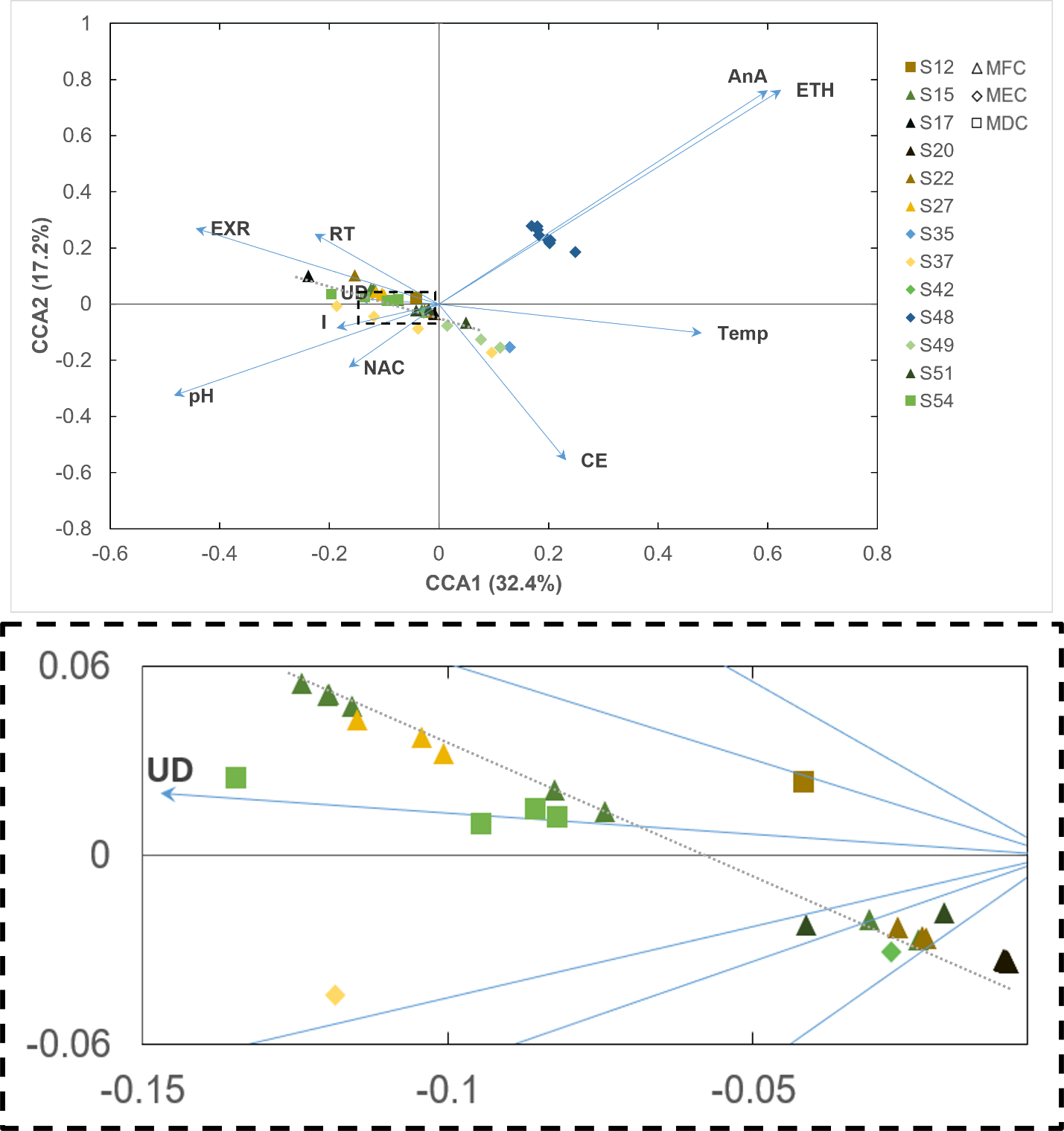

Figure S3. RDA of the 77 samples in the 13 selected publications. The dotted-box region in upper panel is enlarged in the lower panel. MFC: microbial fuel cells, MEC: microbial electrolysis cells, MDC: microbial desalination cells.

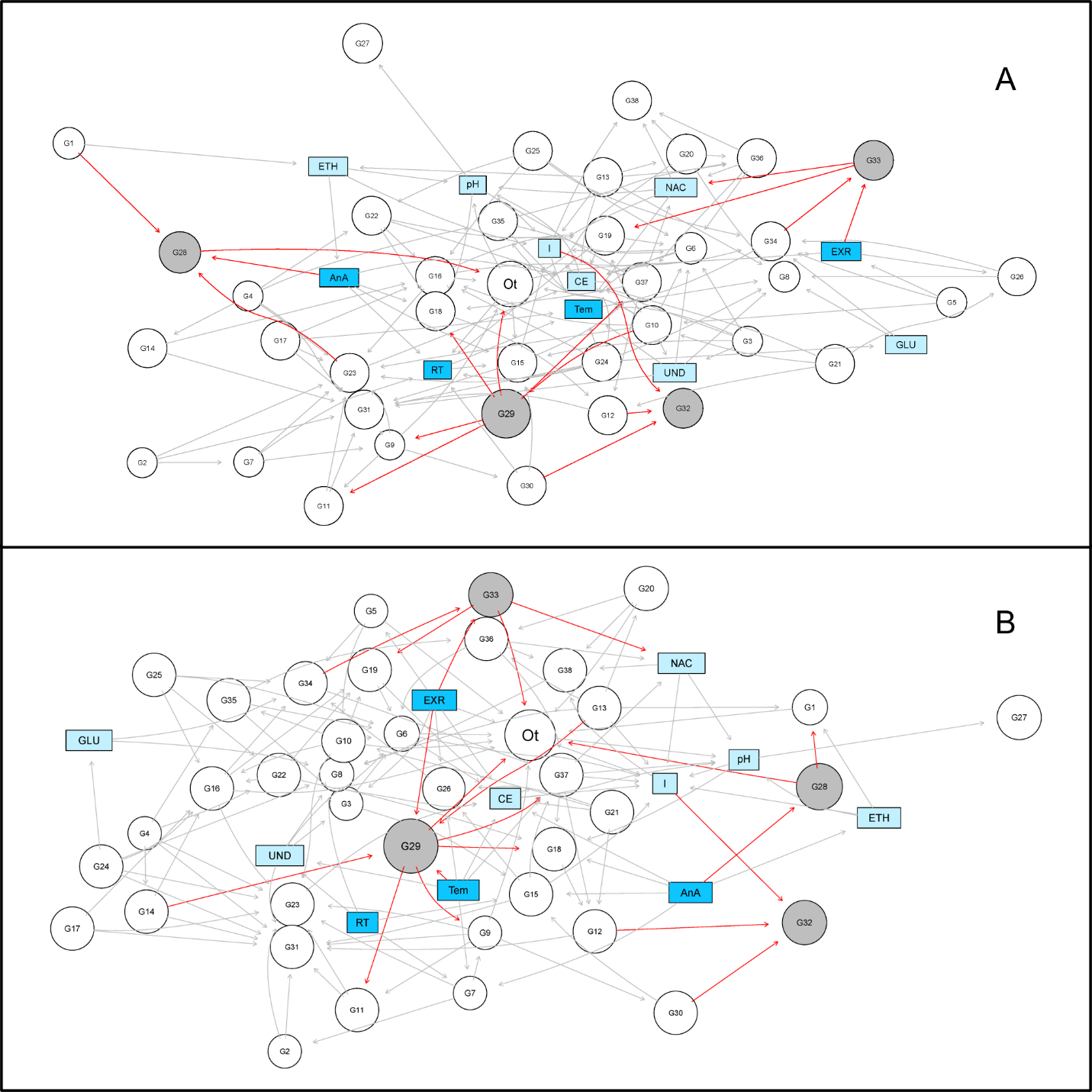

Figure S4. Bayesian networks at the genus level. (A) Primary network: without microbial kinetic parameters input. (B) directed network: without microbial kinetic parameters input and edges pointing to fixed environmental factors.

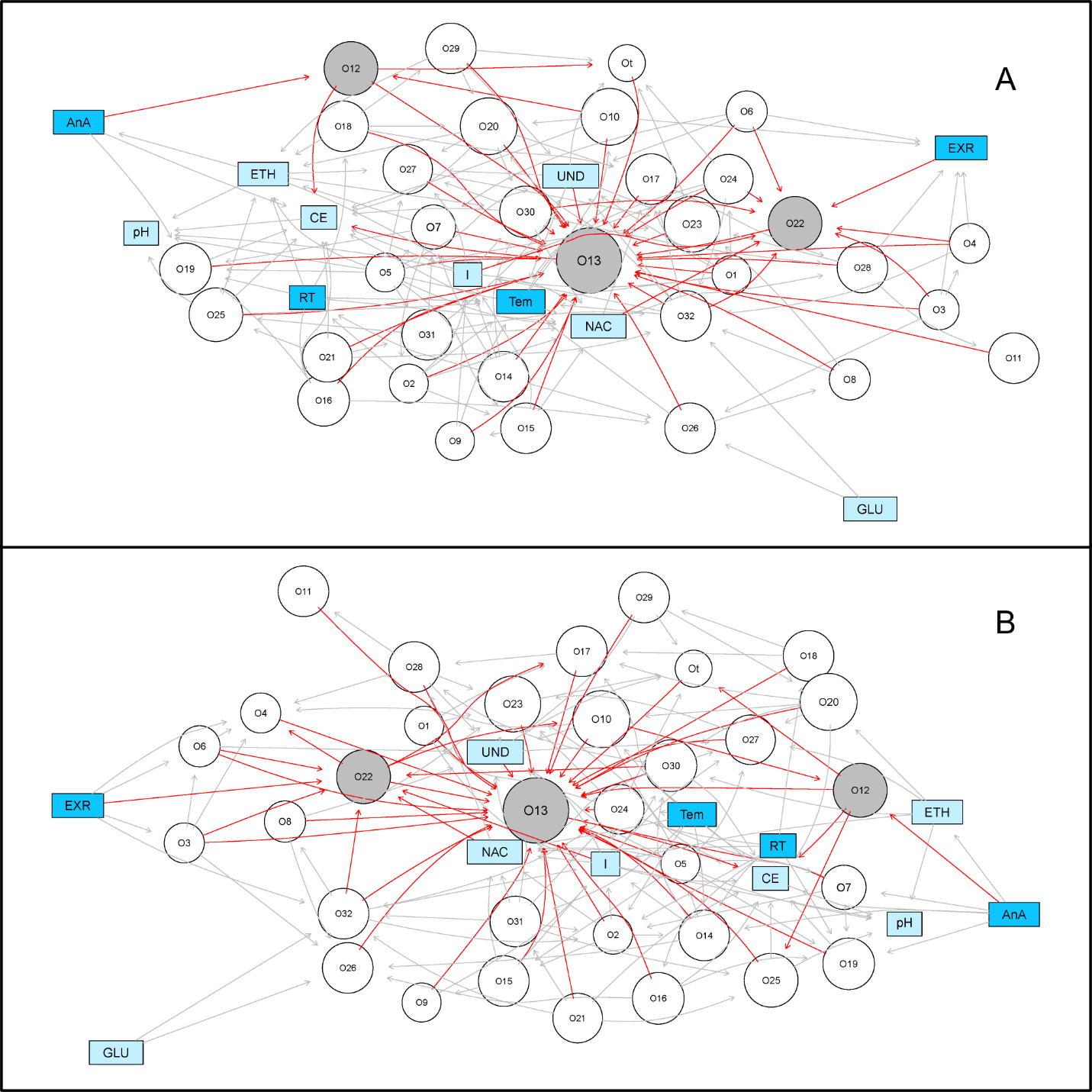

Figure S5. Bayesian networks at the order level. (A) Primary network: without microbial kinetic parameters input. (B) Directed network: without microbial kinetic parameters input and edges pointing to fixed environmental factors.

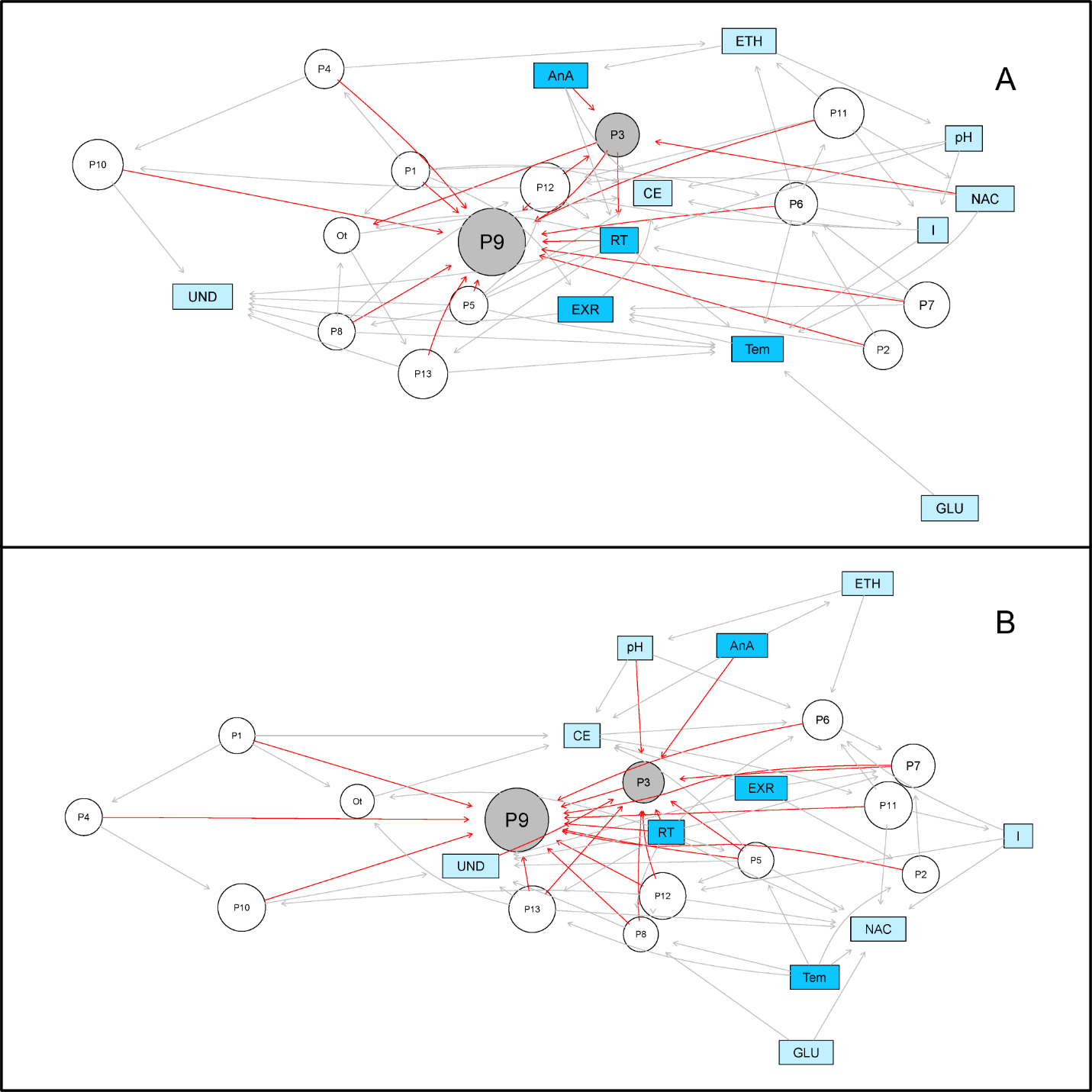

Figure S6. Bayesian network at the phylum level. (A) Primary network: without microbial kinetic parameters input. (B) Directed network: without microbial kinetic parameters input and edges pointing to fixed environmental factors.

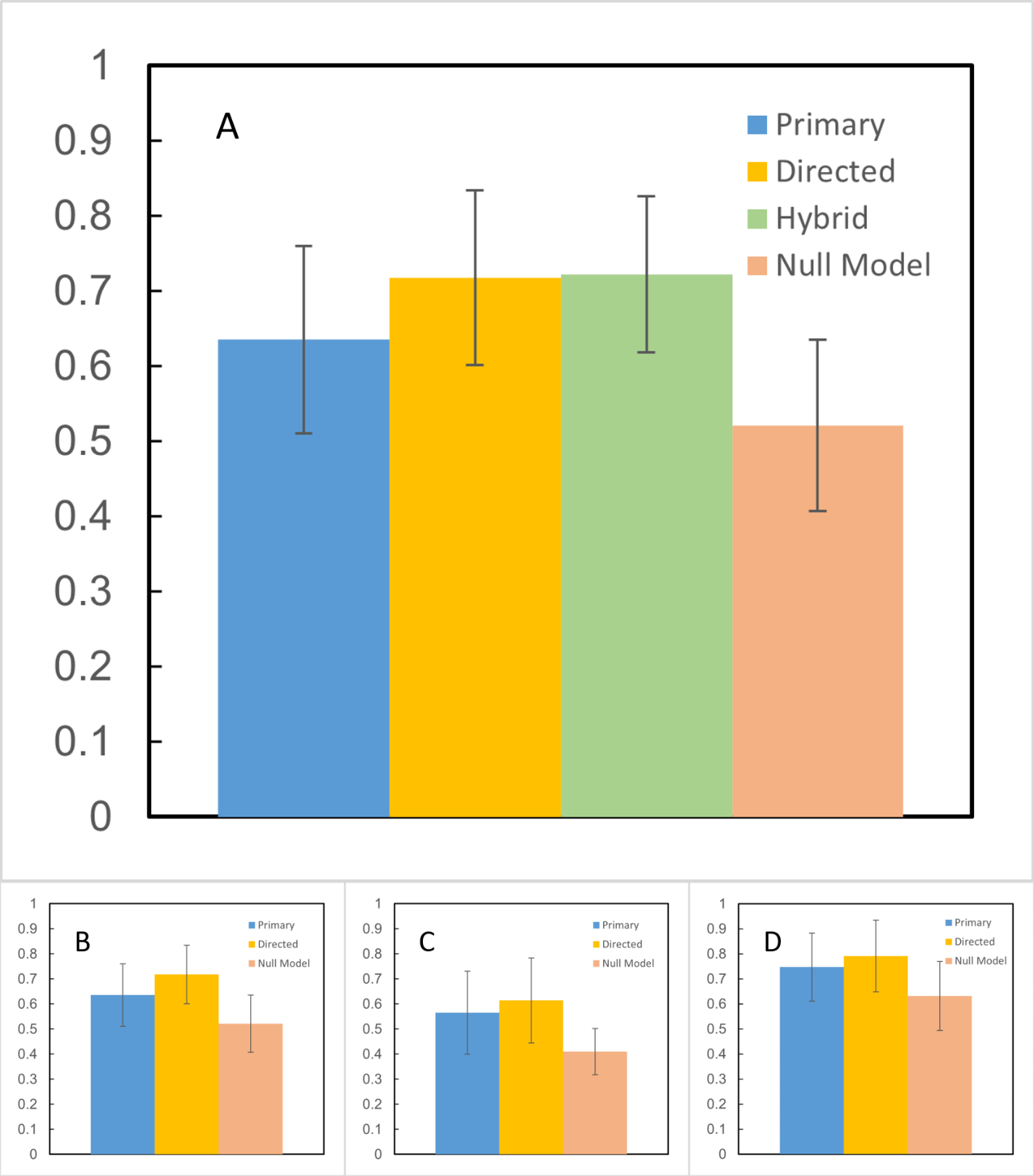

Figure S7. Bray-Curtis similarity between the observed and predicted microbial community from Bayesian networks. Validation at the (A) genus level with hybrid network, (B) genus level without hybrid network, (C) order level, and (D) phylum level.

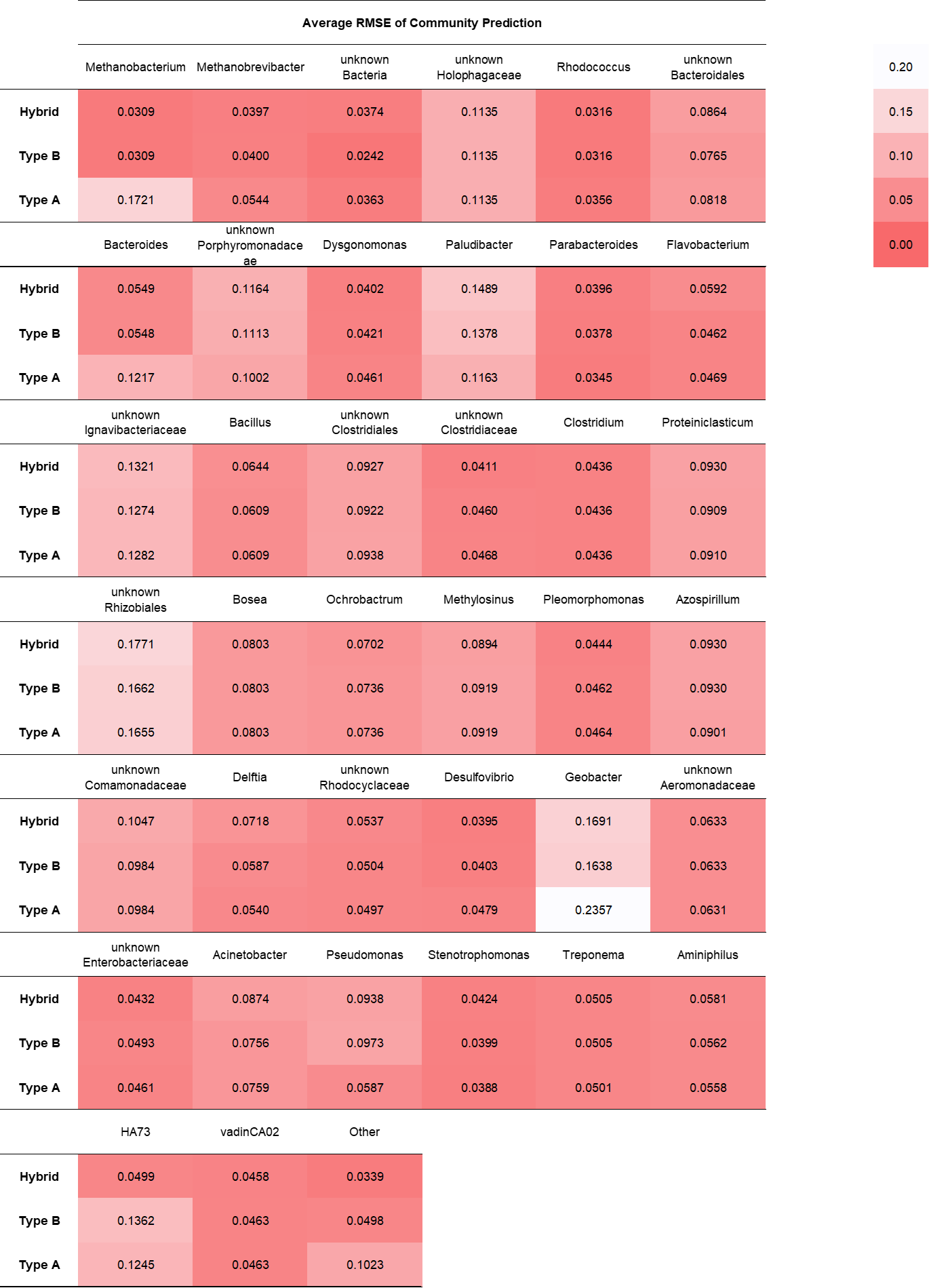
 (Continued)

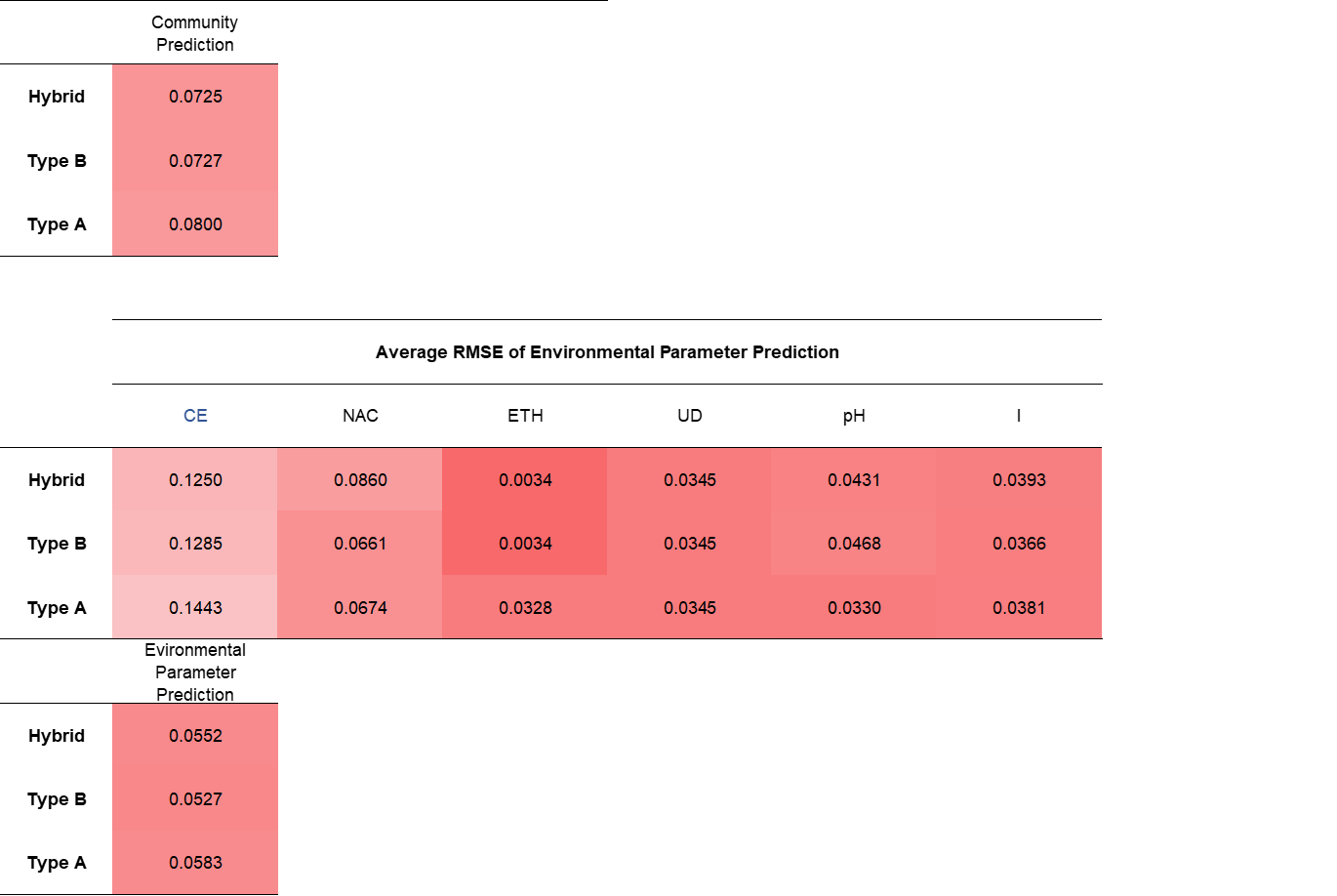

Figure S8. Heat maps of average RMSE at the genus level.

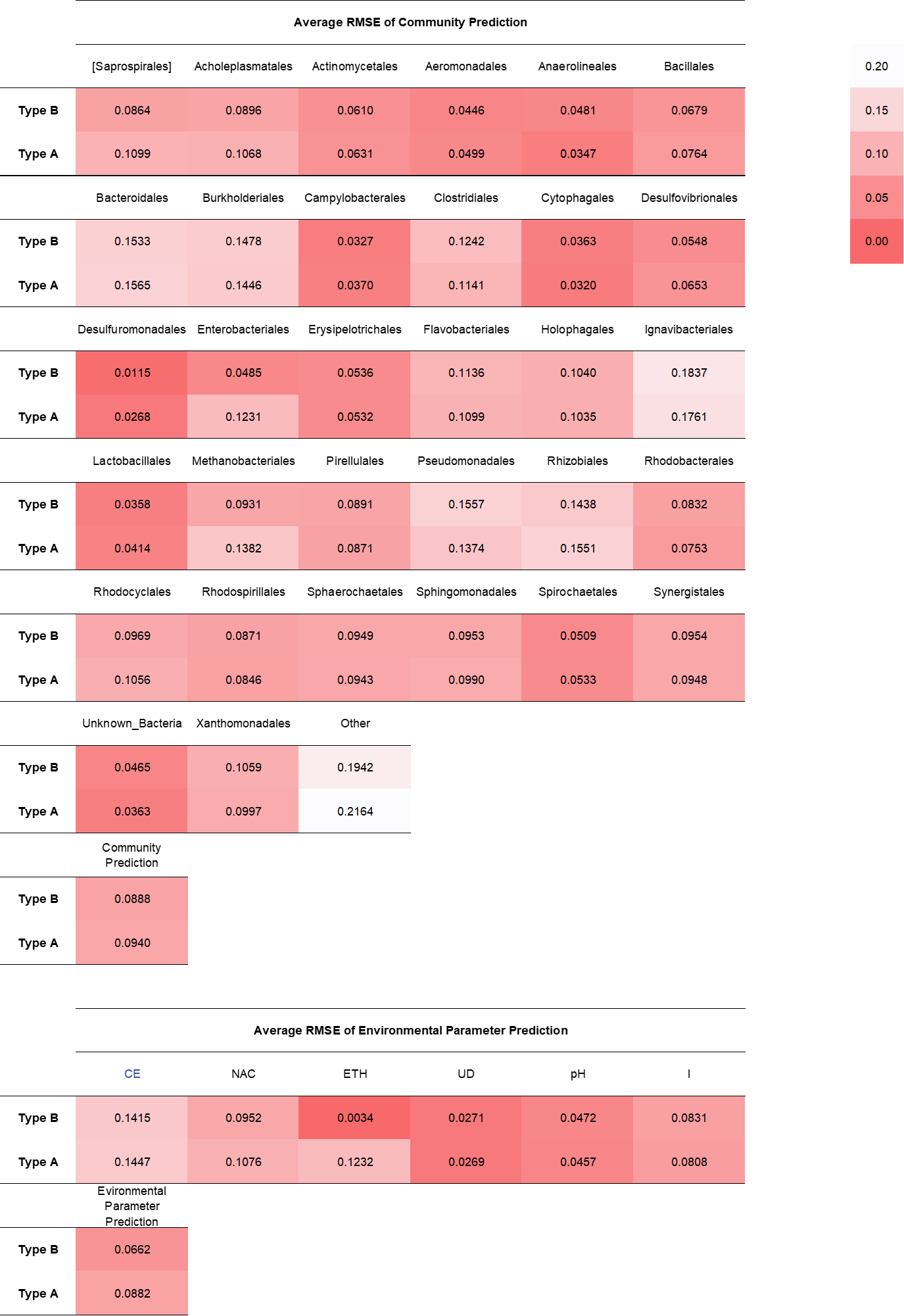

Figure S9. Heat maps of average RMSE at the order level.

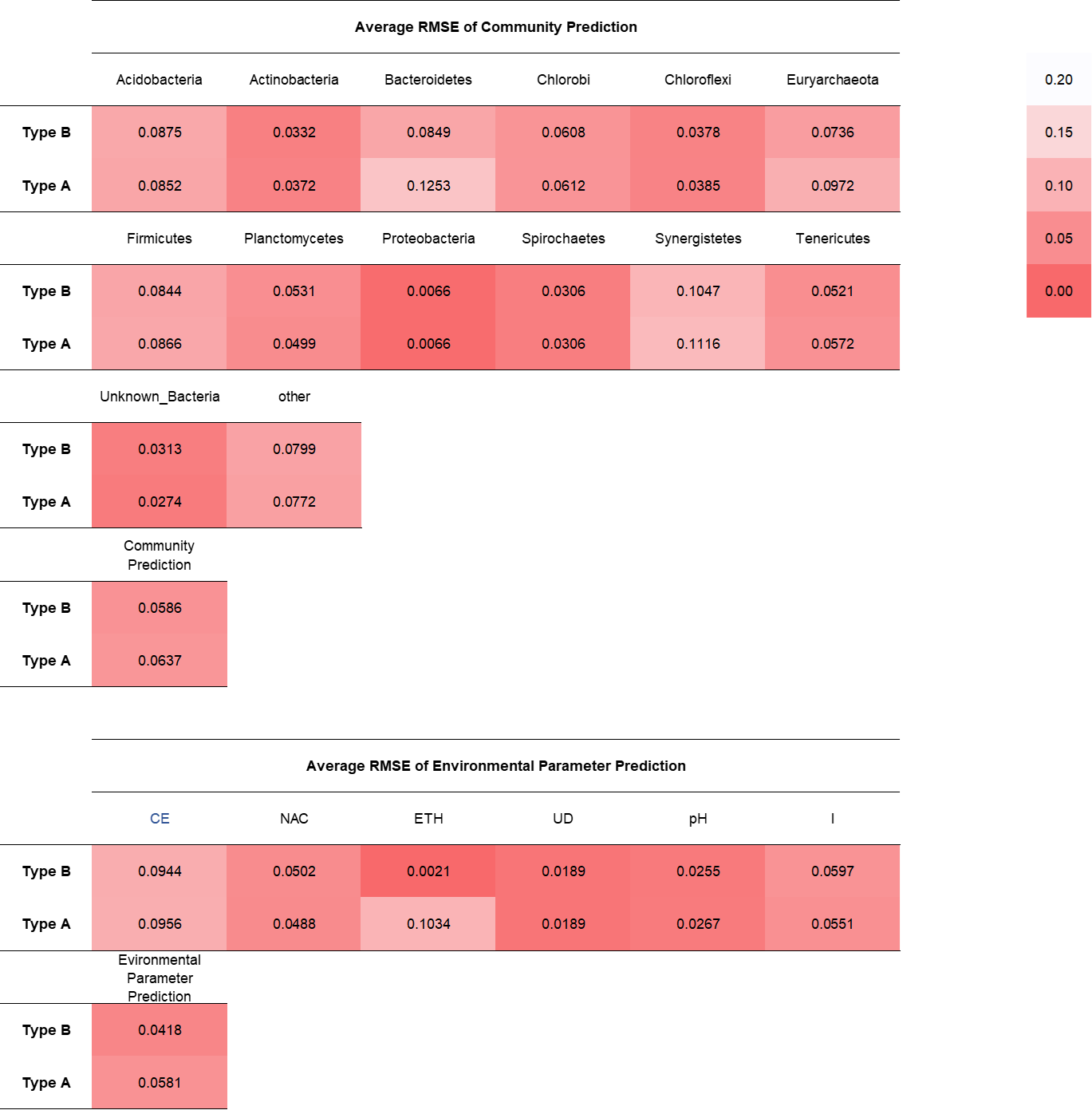

Figure S10. Heat maps of average RMSE at the phylum level.

Table S1. Meta data of the 77 samples from the 13 selected publications.

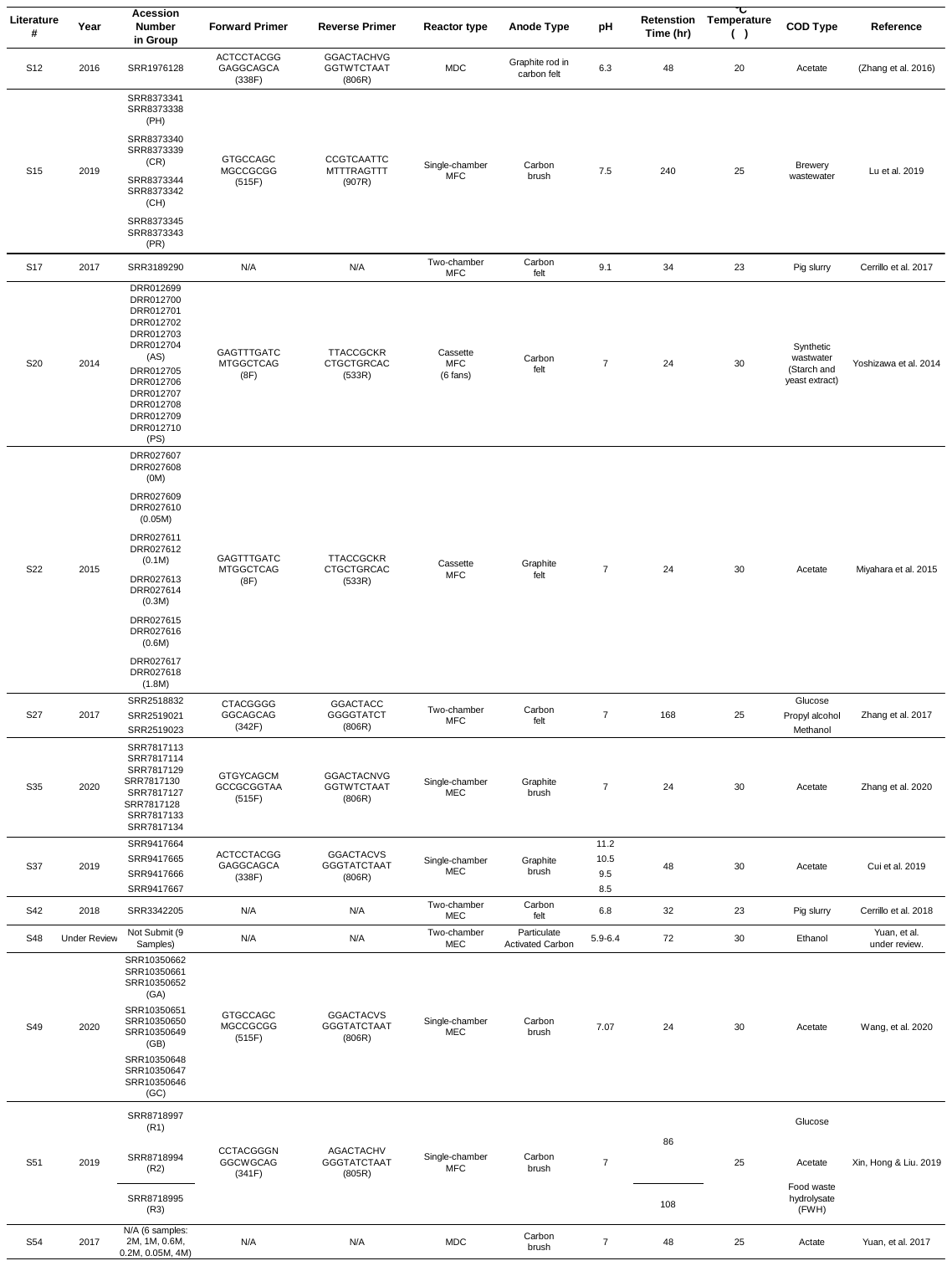

Table S2. The parameters used in the mechanistic component.

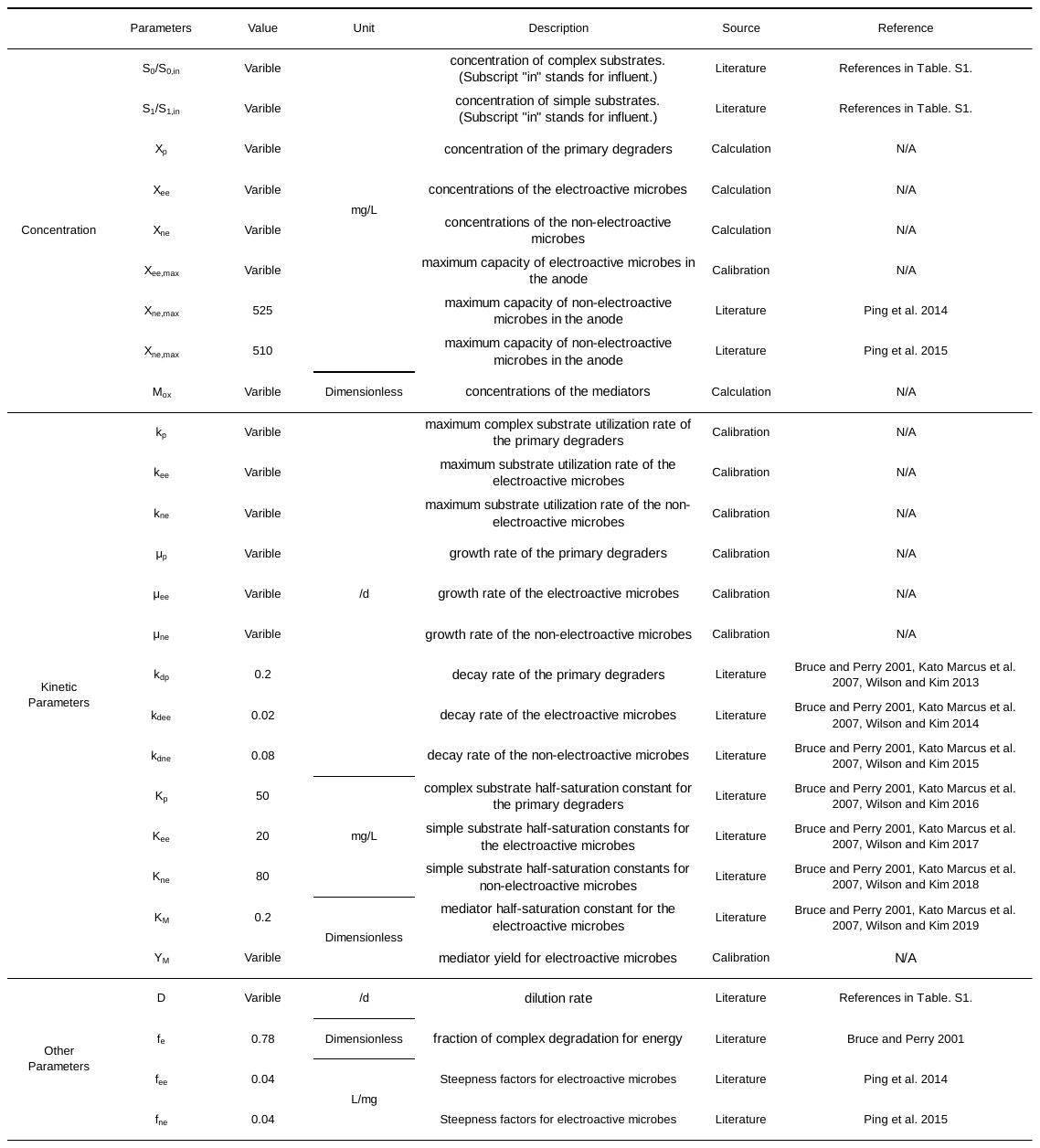

Table S3. Specialized difference equations for calculation of substrate concentration changing over time. To ensure the stability of the difference equations for S20, S22, and S54, $\Delta t$ must be small enough to cancel the increasing vibration of the numerical solution. However, due to the deficiency of the available data, it is unfeasible to obtain stable solutions solely using data from the paper. Interpolation and calibration are the two potential methods to supplement insufficient data. Thus, current data are fitted with a twelve-order polynomial using MATLAB. Then, a time step of 0.864s (e.g., 10^-6^d) was used to calculate the difference equation.

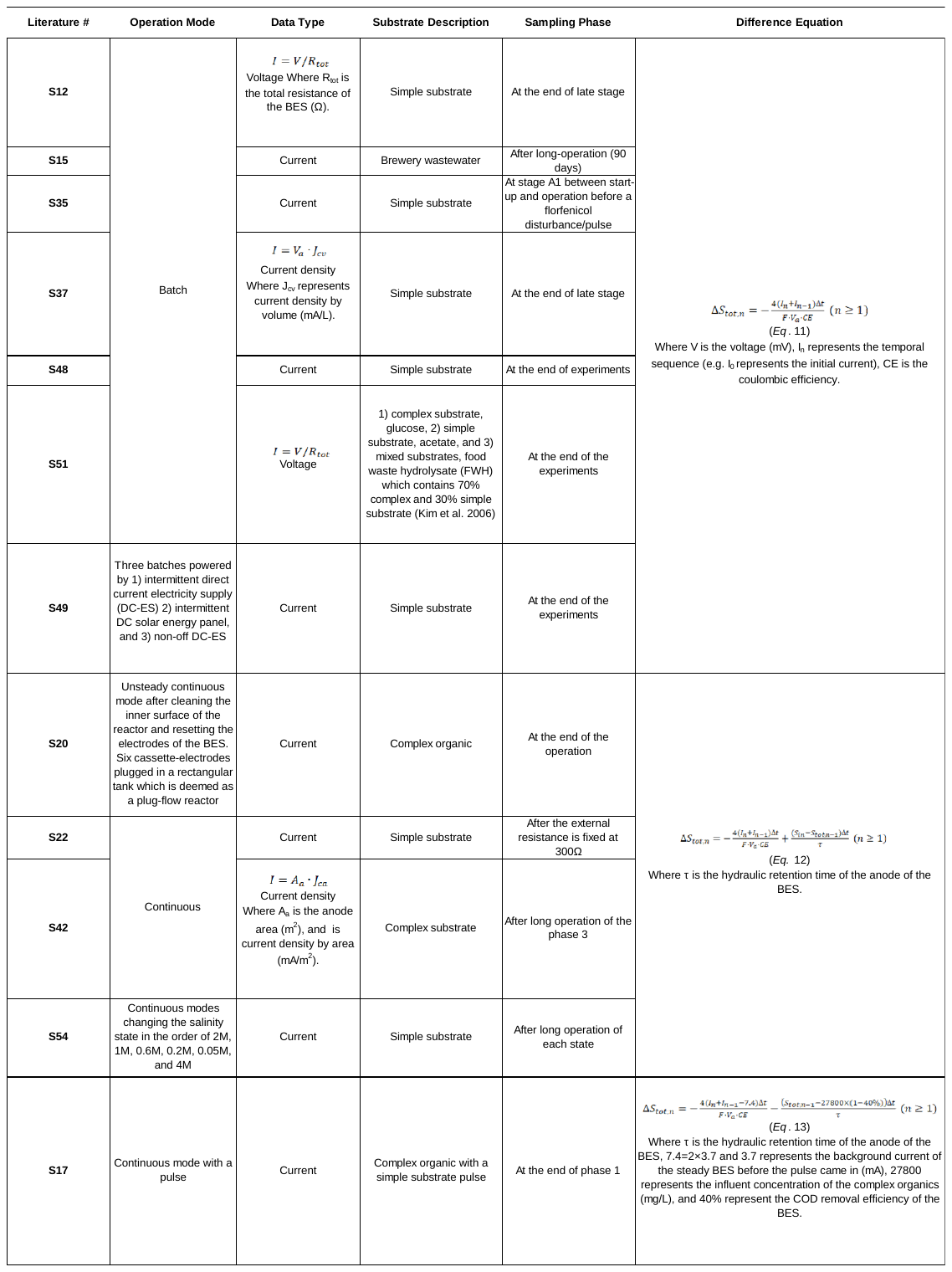

Table S4. Environmental factors and microbial kinetic parameters of the 77 samples for network construction.

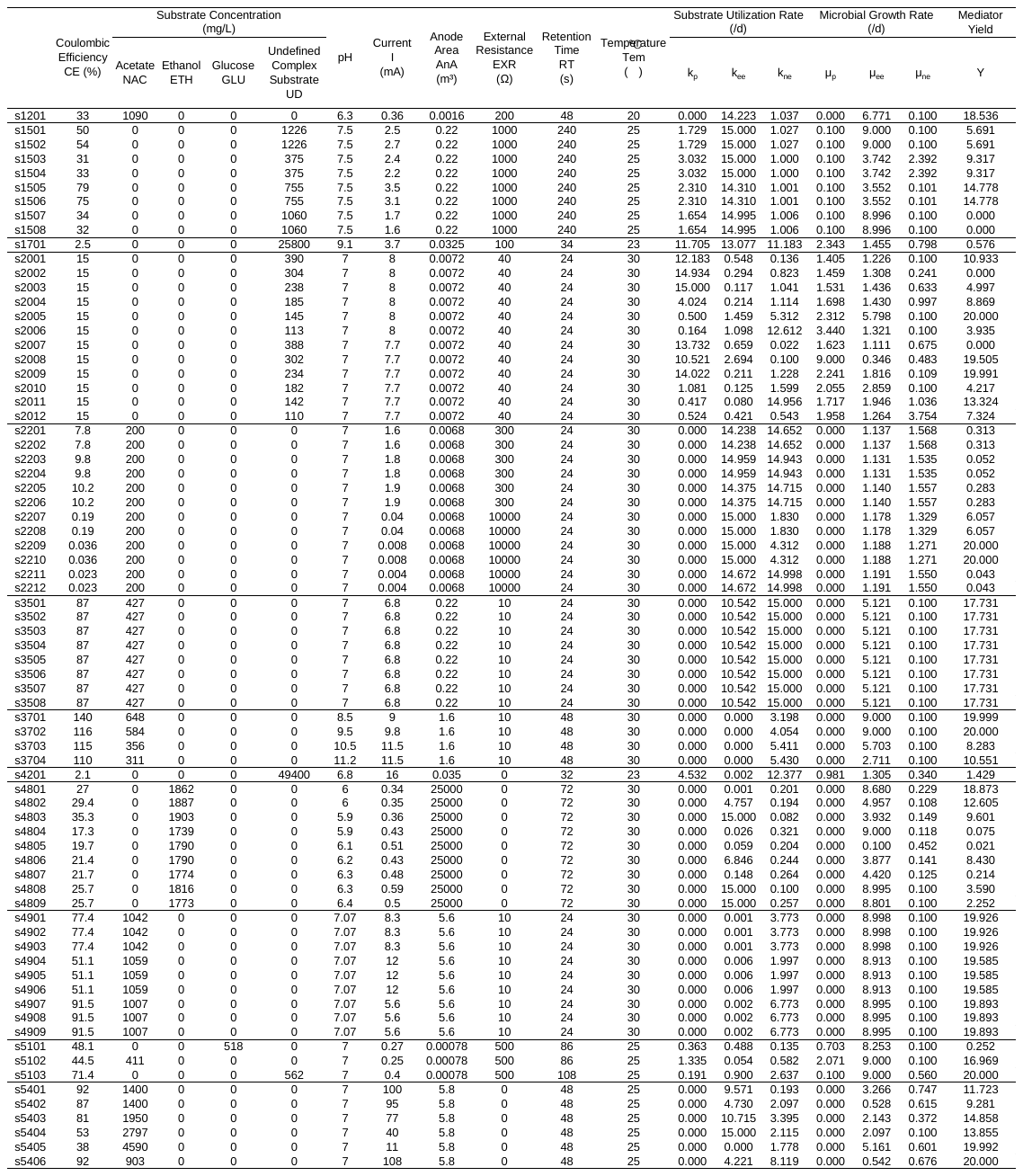

Table S5. Alpha diversity indices.

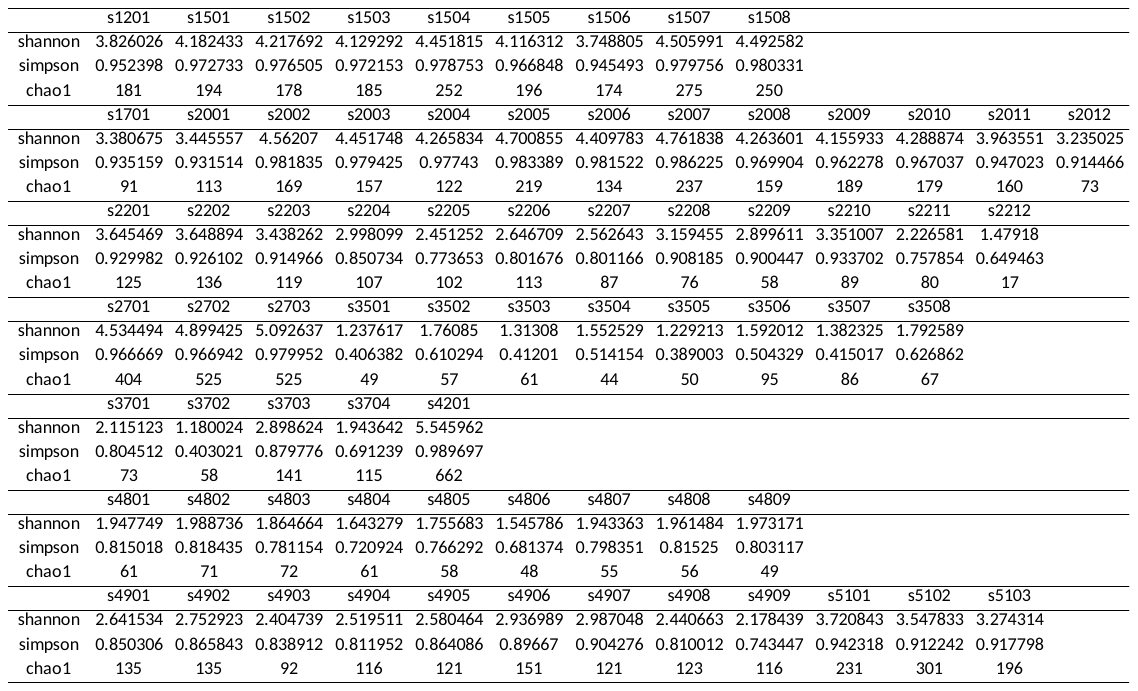

Table S6. Symbols for the core taxa at the genus, order, and phylum levels.

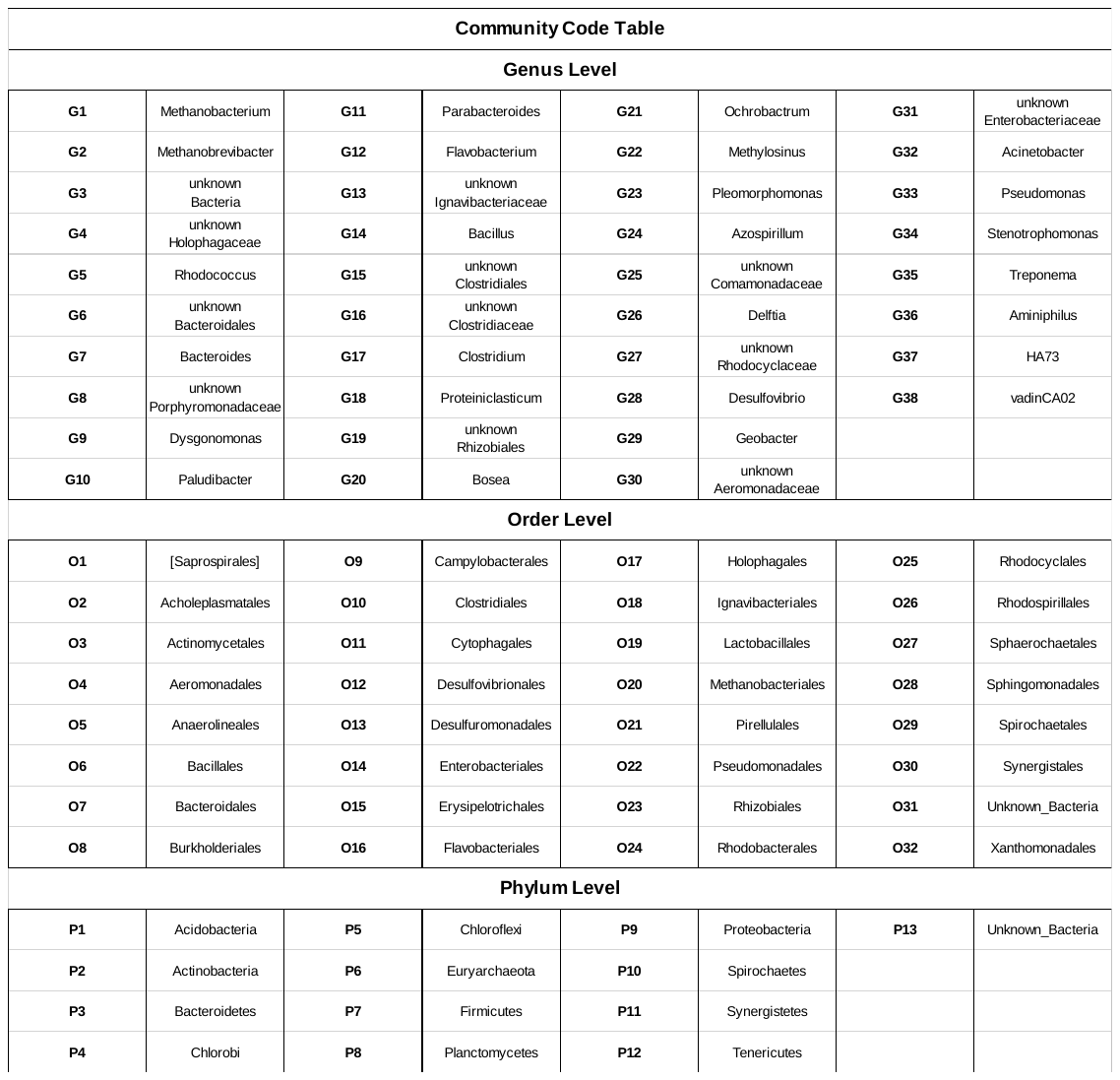

Table S7. Meta data of the six additional samples for testing the robustness of the hybrid model.

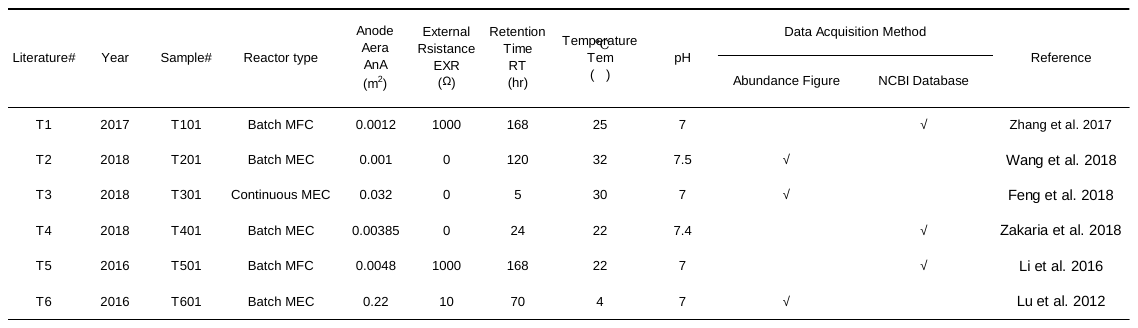
